# Microglial epigenetic memory is associated with accelerated resolution of inflammatory pain induced by prophylactic macrophage-derived small extracellular vesicles

**DOI:** 10.64898/2026.01.15.699551

**Authors:** Xuan Luo, Jason R. Wickman, Jason T. DaCunza, Yuzhen Tian, Ahmet Sacan, Seena K. Ajit

**Affiliations:** Department of Pharmacology & Physiology, Drexel University College of Medicine, 245 North 15th Street, Philadelphia, PA 19102, USA; School of Biomedical Engineering, Science & Health Systems, Drexel University, 3141 Chestnut Street, Philadelphia, PA 19104, USA

## Abstract

Small extracellular vesicles (sEVs) including exosomes play an important role in intercellular communication and can exert immunomodulatory effects in recipient cells. We have shown that a single prophylactic intrathecal injection of sEVs from RAW 264.7 macrophages two weeks prior, promotes faster resolution of mechanical and thermal hypersensitivity in the complete Freund’s adjuvant (CFA) mouse model of inflammatory pain. How this long-term memory develops, and how sEVs regulate immune responses are unknown. Recent studies have shown that priming microglia with inflammatory stimuli can enhance or suppress responses to a delayed secondary insult via epigenetic modifications. We hypothesized that prophylactic intrathecal administration of macrophage-derived sEVs confers accelerated resolution of inflammatory pain by reprogramming epigenetic memory in spinal microglia in recipient CFA model mice. To determine whether prophylactic sEVs could attenuate pain in the absence of microglia when administering sEVs, we ablated microglia using a colony-stimulating factor 1 receptor (CSF1R) inhibitor, PLX5622. sEV-induced pain prophylaxis was completely abolished in PLX5622-fed mice, indicating that microglia are required to be present during sEV administration to confer early resolution of inflammatory pain hypersensitivity. ChIP-seq analysis in spinal microglia 14 days after sEV administration (prior to CFA) revealed an increased number of gene loci enriched for H3K4me1, a hallmark of innate immune memory. Furthermore, inhibiting the H3K4 mono-methyltransferase SETD7 abolished sEV-induced pain attenuation. Our findings indicate that both microglia and its epigenetic reprogramming contribute to pain prophylaxis induced by macrophage-derived sEVs, providing novel insights into the development of non-addictive preventive analgesia.

## Introduction

Neuroimmune interactions in pain engage multiple signaling pathways and cell types, including neurons, glia, and infiltrating immune cells, that together regulate both peripheral and central mechanisms of nociception [1]. These interactions rely on coordinated, cell type and stimulus-specific transcriptional programs involving the dynamic regulation of large gene networks. Transcriptional responses are governed by the accessibility of RNA polymerase and transcription factors to defined genomic regions, a process tightly regulated by chromatin structure through histone modifications and DNA methylation [2, 3]. Inflammatory stimulation can induce epigenetic changes in cells, some of which are rapidly reversible, while others appear to be long-lasting and poised for gene transcription in response to secondary triggers. These persistent epigenetic alterations have been increasingly recognized as a mechanism by which immune cells retain a functional memory of prior inflammatory experiences and linked to innate immune memory [4, 5].

Traditionally, immunological memory was thought to exist only in the adaptive immune system. T and B lymphocytes, major components of the adaptative immunity, express a diverse repertoire of highly specific antigen receptors generated through the rearrangement of the antigen receptor genes, allowing these cells to mount tailored immune responses against various antigens [6]. When naïve, antigen-specific lymphocytes recognize an antigen, they undergo clonal expansion and produce memory cells that can persist for years, ready to respond swiftly and effectively to future exposures of the same antigen. However, recent findings show that innate myeloid and lymphoid cells also retain memory of prior pathogen exposure and become primed to elicit an enhanced response to a secondary stimulus [7, 8]. Unlike the specificity of the adaptive immunity driven by antigen receptors like immunoglobulins and T cell receptors, innate immune cells can provide non-specific responses to homologous (same as the primary encounter) or heterologous (different from the prior stimulus) pathogens or non-microbial ligands. This phenomenon termed innate immune memory or trained immunity is a vital survival mechanism for organisms like plants and invertebrates that lack an adaptive immune system, and for mammals that are deficient in functional T and B cells [9]. Besides trained immunity, innate immune memory can manifest as immune tolerance, when trained cells produce a diminished response to subsequent stimulation. The enhanced or suppressed effects of this response largely depend on the magnitude and duration of the initial stimulus. For instance, microglia treated with a low dose of lipopolysaccharide (LPS), exhibit immune training via increased pro-inflammatory cytokine release upon re-stimulation with a fixed dose of LPS. Conversely, a high initial dose of LPS triggers immune tolerance in microglia, resulting in reduced pro-inflammatory but increased anti-inflammatory cytokine release upon rechallenge [10].

Immune stimulation induces dynamic changes in histone modifications, including histone H3 lysine 4 mono- and tri-methylation (H3K4me1, H3K4me3) and histone H3 lysine 27 acetylation (H3K27ac), leading to lasting reprogramming of the enhancer landscape [11–13]. A hallmark of innate immune memory is the acquisition and persistence of H3K4me1 at enhancers after stimulus withdrawal, even as H3K27ac and transcription factor occupancy are lost, leaving enhancers in a primed state poised for rapid gene induction upon restimulation [12]. In macrophages, these so-called latent enhancers are unmarked at baseline but acquire enhancer features following inflammatory stimulation, retaining H3K4me1 after resolution and enabling accelerated transcriptional responses to secondary challenges [12]. Importantly, similar stimulus-dependent chromatin remodeling has been described in other innate immune cells, including microglia, which can exhibit altered responsiveness to delayed secondary insults through persistent epigenetic modifications [14, 15]. Collectively, these persistent enhancer and promoter changes constitute a form of innate immune memory, commonly referred to as trained immunity [8, 12, 15].

Small extracellular vesicles (sEVs), including exosomes, are key mediators of intercellular communication and can exert potent immunomodulatory effects. However, the mechanisms by which sEVs influence pain hypersensitivity remain poorly understood. Innate immune cell–derived sEVs are known to regulate both innate and adaptive immune responses [16] and accumulating evidence suggests that sEVs can be protective in a range of painful conditions [17–21]. We previously demonstrated that a single prophylactic intrathecal administration of RAW 264.7 macrophage-derived sEVs two weeks prior to injury accelerates the resolution of mechanical allodynia and thermal hyperalgesia in a complete Freund’s adjuvant (CFA) mouse model of inflammatory pain [20].

Notably, the mechanisms by which sEVs confer long-lasting prophylactic effects in naïve mice remain unknown. Microglia, the tissue-resident macrophages of the central nervous system (CNS), play a central role in the development and maintenance of both neuropathic [22] and inflammatory pain [23, 24] through sustained activation and production of pro-inflammatory cytokines, including IL-1β, IL-6, and TNF-α [25, 26]. We hypothesized that prophylactic intrathecally administered RAW 264.7 macrophage-derived sEVs confer faster resolution of CFA-induced pain hypersensitivity by reprogramming epigenetic memory in spinal microglia of the recipient mice.

## Materials and methods

### Cell culture

RAW 264.7 macrophage cells (#TIB-71, American Type Culture Collection) were cultured in complete Dulbecco’s Modified Eagle’s Medium (DMEM; Corning, New York, NY) supplemented with 10% heat-inactivated fetal bovine serum (FBS; #35-011-CV, Corning) and 1% penicillin-streptomycin (P/S; ThermoFisher Scientific, Waltham, MA) in a humidified incubator at 37°C with 5% CO_2_.

### Preparation of exosome-depleted FBS

Heat-inactivated FBS was added to 70-mL ultracentrifuge tubes (#355622, Beckman Coulter, Brea, CA) and centrifuged in an Optima LE-80K ultracentrifuge (Beckman Coulter) with a Type 45 Ti fixed-angle rotor (Beckman Coulter) at 120,000xg for 18 h at 4°C to remove larger vesicles and aggregates, including exosomes. The supernatant was filtered through a 0.22µm syringe filter and stored at −20°C for further use.

### Isolation of small extracellular vesicles (sEVs) from conditioned medium

To obtain sEV-conditioned medium, RAW 264.7 cells were plated in T-175 flasks (Corning) at a density of 29,000 cells/cm^2^. Upon reaching 70-80% confluency, cells were washed three times with 10 mL of phosphate-buffered saline (PBS; Corning) each, followed by replacement with 35 mL of DMEM containing 10% exosome-depleted FBS and 1% P/S. Following a 24-h incubation period, conditioned medium was collected in a 50-mL conical tube and centrifuged at 500×*g* for 10 min at 4°C. The resulting supernatant was transferred to a new conical tube and stored at −80°C until further processing. To enhance both the recovery and purity of sEV preparation, we followed the Minimal Information for Studies of Extracellular Vesicles (MISEV) guidelines [27] and employed a multi-step approach involving filter concentration, size-exclusion chromatography (SEC), and differential ultracentrifugation. All centrifugations were performed at 4°C. Thawed media was centrifuged at 12,000xg for 35 min (Sorvall RC-5C Plus centrifuge with Sorvall SA-600 fixed-angle rotor, ThermoFisher Scientific), and the supernatant was passed through a 0.22μm syringe filter. The supernatant was then concentrated to less than 500 µL using 100 kDa Amicon centrifugal filters (#UFC9100, Millipore, Burlington, MA) through centrifugation at 5,000xg for 35 min each. The retentate was diluted to 500 µL with PBS and subjected to purification by SEC using a qEV original 35 nm Legacy column (#SP5, Izon Science, New Zealand) according to the manufacturer’s instructions. Four EV-rich fractions (7-10, 0.5 mL each) were combined, followed by an ultracentrifugation at 110,000xg for 70 min (Optima TLX ultracentrifuge with TLA 100.4 rotor, Beckman Coulter). The final sEV pellet was resuspended in 35 µL of PBS and stored at −80 °C. The protein concentration of each sEV preparation was quantified using the microassay protocol for the DC protein assay (Bio-Rad, Hercules, CA).

### Nanoparticle tracking analysis (NTA) of sEVs

The size distribution and concentration of sEVs were measured by tracking their Brownian motion under laser illumination using a NanoSight LM10 instrument equipped with a 405 nm laser (Malvern Panalytical). Sample diluent (PBS) was filtered through 0.1µm filters before use. Samples were injected into the sample chamber with sterile syringes and further diluted until an optimal concentration of 20-90 particles/frame was achieved. Five 60-second videos were captured for each sample. The temperature during sample acquisition was also recorded for analysis. Samples were diluted to a particle concentration of roughly 1×10^8^/mL and measured using the ZetaView S/N 18-390 (ParticleMetrix) and analyzed using the ZetaView software (version 8.05.12 SP2).

### Western blot for sEV markers

Both RAW 264.7 cells and sEVs were thawed and lysed in radioimmunoprecipitation assay (RIPA) buffer (ThermoFisher Scientific) with 1X protease inhibitor cocktail (ThermoFisher Scientific). Protein concentration for each cell sample was measured using the DC protein assay. Equal amounts of protein (5 μg) from cell lysates and sEVs were resolved on 4-12% gradient SDS-PAGE gel (ThermoFisher Scientific) and transferred to a 0.45-μm PVDF membrane. Membranes were blocked with 5% non-fat dry milk (NFDM) in Tris-buffered saline with 0.1% Tween 20 (TBST) for 1 hour at room temperature on an orbital shaker. After washing 3x in TBST, membranes were incubated with primary antibodies diluted in TBST + 5% NFDM overnight at 4°C on an orbital shaker. Membranes were washed 3x in TBST to remove excess primary antibody and incubated with secondary antibodies in TBST + 5% NFDM at room temperature for 1 h on an orbital shaker. Membranes probed with HRP-conjugated antibodies were incubated with SuperSignal West Dura Substrate (ThermoFisher Scientific) for 5 min at room temperature. Membranes were visualized using the Odyssey Fc imaging system (LI-COR Biosciences). The following primary antibodies and concentrations were used: mouse anti-CD81 Clone B-11 (1:500, #sc-166029, Santa Cruz Biotechnology), mouse anti-Alix Clone 1A12 (1:300, #sc-53540, Santa Cruz Biotechnology), and rabbit anti-Calnexin (1:5,000, #PA5-34754, Santa Cruz Biotechnology). Secondary HRP-conjugated antibodies used were Goat anti-rabbit IgG-HRP and Goat anti-mouse IgG-HRP (#1706515 and #1706516, Bio-Rad).

### Detection of sEV surface epitopes by flow cytometry

The mouse MACSPlex EV Kit (#130-122-211, Miltenyi Biotec, Germany) was used to detect 37 immune markers using BD LSRFortessa flow cytometer (BD Biosciences, San Diego, CA) following the manufacturer’s protocol using 10 µg of sEVs as previously described [28]. sEVs are identified by CD9^+^, CD63^+^, and CD81^+^ APC-labelled capture beads, which are then labelled with marker specific antibodies. For markers with %Pos ≥ 60%, the weighted expression is calculated as: 0.5 * %Pos * (1 + MFI / Highest MFI). Samples are represented in the form of a heat map. Three individual sEV preparations were used.

### Mice

Wild-type (WT) C57BL/6J male mice aged 6-9 weeks were purchased from the Jackson Laboratory (Bar Harbor, ME) and housed socially (2-5 mice per cage) in individually ventilated cages in a temperature- and humidity-controlled room with a 12-h light/dark cycle. Mice were provided with irradiated rodent diet (#5053, LabDiet, St. Louis, MO) and water *ad libitum* unless otherwise specified. Upon arrival, mice were acclimated in the animal facility for at least one week prior to experiments. All procedures involving animals were carried out in accordance with the Guide for the Care and Use of Laboratory Animals of the National Institutes of Health and approved by the Institutional Animal Care & Use Committee of Drexel University College of Medicine.

### Intrathecal injection of sEVs

Mice received intrathecal injections at 8-11 weeks of age. A Hamilton microliter syringe equipped with a 30-gauge needle was inserted between the L5 and L6 vertebrae at a 30° angle. Successful entry into the intradural space was confirmed by observing a tail flick response. One µg of sEVs resuspended in 10 µL of PBS (0.1 µg/µL) or an equal volume of PBS, was injected slowly and steadily.

### Complete Freund’s adjuvant (CFA) model of inflammatory pain

To induce inflammatory pain in mice, 20 µL of 50% CFA (10 µg, #F5881, Sigma-Aldrich, St. Louis, MO) emulsion in 0.9% saline was administered subcutaneously into the plantar surface of the right hind paw using a 28-gauge needle and syringe. In the prophylactic pain model, mice received an injection of CFA two weeks after a single dose of 1 µg sEVs [20].

### Pain behavioral assessment of mechanical hypersensitivity

Mechanical allodynia was assessed using the von Frey test on mouse hind paws, as previously described [20]. Mice were habituated by being individually placed into plexiglass chambers on an elevated mesh grid platform for the von Frey test for 1 h, 1–2 days prior to baseline testing. A 30-min habituation period was also conducted on each day of behavioral testing. To test mechanical sensitivity, a series of von Frey filaments (#BIO-VF-M, Bioseb, France) ranging in ascending order from 0.02 to 1.4 grams of force were sequentially applied to the center of the paw plantar surface. A positive response to the poking was recorded when mice withdrew, licked, or flinched the testing paw. Each filament was applied five times for 2-3 sec each. The paw withdrawal threshold was determined as the force at which three positive responses out of five stimuli were observed. Measurement of sensitivity in unaffected paws (naïve mice or the contralateral (left) paw of CFA model mice) began with a 0.16 g filament, while the ipsilateral (right) paw of CFA model began with a 0.02 g filament. The experimenter was blinded to the treatment conditions.

### PLX5622-mediated microglia depletion

Mice were fed control AIN-76A rodent diet (#D10001i, Research Diets, New Brunswick, NJ) *ad libitum* upon arrival from the Jackson Laboratory. To pharmacologically ablate microglia in the spinal cord, a CSF1R inhibitor, PLX5622 (#HY-114153, MedChemExpress, Monmouth Junction, NJ), was formulated in AIN-76A chow containing 1,200 parts per million (ppm) of PLX5622 (#D19101002i, Research Diets). PLX5622 diet was fed to 8-9 week-old mice starting 10 days before the administration of sEVs for a duration of 14 days. These mice were reverted to the control diet for the remainder of the study period.

### Intrathecal delivery of drugs

To enable repeated drug administration into the intrathecal space of mice, a polyurethane intrathecal catheter (32 G tip, 6 cm long, 22 G connection; #MIT-02, SAI Infusion, Lake Villa, IL) was inserted between the L5 and L6 vertebrae into the subarachnoid space and extended to lumbar L4 to L5 regions as previously reported [29]. The catheter was secured with 4-0 sutures under the skin, and 7 µL of PBS was injected into the catheter to flush, followed by heat sealing of the tube opening. Mice were allowed to recover for 5–7 days before experimentation. PFI-2 (#S7294, Selleck Chemicals, Houston, TX), a specific inhibitor of SET domain-containing 7 histone lysine methyltransferase (SETD7), was used to suppress the methyltransferase activity of SETD7. PFI-2 was reconstituted in dimethyl sulfoxide (DMSO; G-Biosciences, St. Louis, MO) at a concentration of 10 mM and stored at −20°C. The injection regimen was as follows. Day 1: 5 µL of 20 µM PFI-2 in 1% DMSO, 3 µL of PBS, 5 µL of sEVs containing 1 µg of sEVs (0.2 µg/µL), and 7 µL of PBS. Day 2–4: 10 µL of 10 µM PFI-2 in 0.1% DMSO, followed by 7 µL of PBS. PBS was administered between drugs to avoid direct interaction between PFI-2 and sEVs. Furthermore, to ensure complete delivery of the drugs into the cerebrospinal fluid circulation, PBS was also injected at the end. DMSO and 5 µL of PBS were used as controls for PFI-2 and sEVs, respectively. The catheter was heat-sealed after injections each day. Mice were anesthetized with 2% isoflurane gas during catheterization and injections. Additionally, mice were housed individually until the completion of intrathecal drug delivery to prevent catheter displacement.

### Perfusion, tissue fixation, and immunohistochemistry (IHC)

Mice anesthetized via intraperitoneal (i.p.) injection of 100 mg/kg body weight of ketamine and 10 mg/kg body weight of xylazine were transcardially perfused with room temperature PBS, followed by ice-cold 4% (w/v) paraformaldehyde (PFA; Electron Microscopy Sciences, Hatfield, PA) in 0.1 M phosphate buffer (PB) for 5 min. L4-L5 lumbar spinal cord segments were dissected and post-fixed in 4% PFA in PB overnight at 4°C. Following cryoprotection in 30% sucrose in PB for at least 24 h, tissues were embedded in O.C.T. compound and sectioned at 25 µm in PB (free-floating) at −20°C using a cryostat. For immunostaining, sections were washed three times with wash buffer (0.3% Triton X-100 in PB) for 5 min each, then blocked in wash buffer containing 5% normal goat serum (NGS; #S-1000, Vector Laboratories) for 2 h at room temperature. Sections were incubated with primary antibodies diluted in blocking buffer on a shaker overnight at 4°C. The next day, sections were washed three times with wash buffer, then incubated with secondary antibodies in PB supplemented with 5% NGS for 2 h at room temperature on a shaker. Sections were washed three times with wash buffer and counterstained with 1 µg/mL DAPI (4’,6-diamidino-2-phenylindole; ThermoFisher Scientific) for 10 min at room temperature. Following a final wash step, tissue sections were mounted on clean Superfrost Plus Gold slides and cured with mountant (Invitrogen) overnight in the dark. Confocal images were acquired using an Olympus FV3000 microscope. Antibodies were used as follows: Iba1 (1:2000, #019-19741, Wako Chemicals) and goat anti-rabbit Alexa Fluor 594 (1:1000, #A-11012, Invitrogen).

### Preparation of single-cell suspension from adult mouse spinal cord

Spinal cords from PBS-perfused mice were hydraulically extruded into ice-cold Dulbecco’s Phosphate-Buffered Saline (DPBS) supplemented with calcium and magnesium (DPBS++; #14287072, Gibco) using a trimmed 200-µL pipette tip and a 10-mL syringe filled with cold DPBS++. Tissue dissociation was carried out using the Adult Brain Dissociation Kit for mouse (#130-107-677, Miltenyi Biotec). In brief, spinal cord was minced into 2 mm^2^ pieces and placed in a gentleMACS C tube (Miltenyi Biotec) containing Enzymes P and A. The C tube was then placed on a gentleMACS Octo Dissociator with Heaters (Miltenyi Biotec) and subjected to appropriate programs based on tissue weight (20–100 mg: 37C_ABDK_02; >100 mg: 37C_ABDK_01). Dissociated cells were then passed through a 70-µm cell strainer into a 50-mL conical tube and centrifuged at 300xg for 10 min at 4 °C. Cells were resuspended in an appropriate buffer for downstream experiments.

### Isolation of spinal cord microglia using magnetic-activated cell sorting (MACS)

Single cells from adult mouse spinal cord were resuspended in 3.1 mL of DPBS++ and the suspension was then transferred to a 15-mL conical tube. To remove debris such as myelin, 0.9 mL of debris removal solution provided by the dissociation kit was thoroughly mixed with the suspension. Subsequently, 4 mL of DPBS++ was carefully overlaid on the cell suspension, followed by centrifugation at 3,000xg for 10 min at 4°C. The top two phases were aspirated, and the remaining suspension was diluted to 15 mL with DPBS++ followed by centrifugation at 1,000xg for 10 min at 4°C. Debris-free cells were incubated with CD11b MicroBeads (10 µL beads per 10^7^ cell; #130-049-601, Miltenyi Biotec) in PBS + 0.5% bovine serum albumin (BSA) for 15 min in a refrigerator (2−8 °C). Unbound antibody was removed by adding 1-2 mL of 0.5% BSA in PBS per 10L cells and centrifuging at 300xg for 10 min at 4°C. The magnetic-labeled positive fraction (CD11b^+^ cells) was isolated using an LS column (Miltenyi Biotec). CD11b^+^ cells were resuspended in a suitable buffer for subsequent experiments. Each MACS isolation utilized a pool of 6-8 mice. For RNA isolation, final cell pellets were dissolved in 300 µL of RNA Lysis/Binding buffer and snap-frozen in liquid nitrogen.

### Percoll density gradients

Isolation of mononuclear cells from spinal cord using Percoll gradients was performed as described [30]. First, a stock isotonic Percoll (SIP) was prepared by mixing 9 parts of Percoll (Cytiva, Marlborough, MA) with one part of 10X HBSS without calcium, magnesium, or phenol red (Gibco). SIP was diluted to 30% with complete RPMI 1640 medium (Gibco) containing phenol red, 10% FBS and 1% P/S, and diluted to 70% with 1x HBSS (Avantor, Radnor, PA). After dissociation, cells were resuspended in 10 mL of 30% SIP, transferred to a 15-mL conical tube, and underlaid with 2.5 mL of 70% SIP. Gradients were centrifuged at 500×*g* for 30 min at 18°C with no brake. The supernatant containing the myelin was discarded. Subsequently, 4 mL of cell suspension was collected at the 30%/70% interphase into a new 15-mL conical tube containing 8 mL of 1X HBSS and centrifuged at 500×*g* for 7 min at 18°C. Cell pellets were then resuspended in an appropriate buffer.

### Isolation of spinal microglia using fluorescence-activated cell sorting (FACS)

For isolating microglial cells using FACS, each sample was comprised of a pool of 6-8 mice (n=2). Spinal mononuclear cells isolated using Percoll gradients were washed in FACS buffer (PBS containing 2% FBS, 1% P/S, 25 mM HEPES (Gibco), and 2.5 mM EDTA (Invitrogen)). After buffer exchange, cells were resuspended in 50 µL FACS buffer containing FcR block (1:100; #156603, BioLegend) for 5 min at room temperature, followed by 50 µL of live/dead staining in PBS (1:500; #L10119, Invitrogen) for 15 min at 4°C. Cells were washed and resuspended in FACS buffer. Cells were then co-stained with 55 µL of antibody cocktail in stain buffer (#566349, BD Biosciences) for 45 min at 4°C. Cells were washed and incubated with 100 µL of Brilliant Violet (BV) 785 Streptavidin in FACS buffer (1:100; #405249, BioLegend) for 5 min at room temperature to label biotinylated antibodies (CD3, CD19, and NK1.1) in a dump channel to exclude T cells, B cells, and natural killer (NK) cells. Following a final wash, cells were resuspended in 300 µL of FACS buffer. For each washing step, samples were filled up to 1.2 mL with FACS buffer and centrifuged at 1,500 rpm for 5 min at 4°C. Finally, cells were filtered through a 35-µm cell strainer (#352235, Falcon) immediately before sorting on a Cytek Aurora CS cell sorter (63.6 psi sheath pressure; Cytek Biosciences, Fremont, CA) with SpectroFlo software. All procedures were conducted with minimal light exposure. Primary antibodies were used as follows: BUV395 CD45 (1:50; #564279, BD Biosciences), APC CD11b (1:50; #561690, BD Biosciences), PE/Dazzle 594 Ly6C (1:50; #128043, BioLegend), PerCP/Cyanine5.5 Ly6G (3:50; #127615, BioLegend), biotin CD3 (3:50; #100243, BioLegend), biotin CD19 (1:25; #115504, BioLegend), and biotin NK1.1 (1:50; #108703, BioLegend). Microglia were sorted as CD45^+^CD11b^+^Ly6C^-^Ly6G^-^Dump. For each collection of microglia from CFA model mice, ipsilateral lumbar (L1–L6) spinal cords were pooled from 6-8 mice.

### Tissue homogenization and RNA isolation

Dissected L4-L5 spinal cord was snap-frozen in 1.5-mL nuclease-free tubes in dry ice or liquid nitrogen and stored at −80°C. Immediately before disruption, 3–4 ceramic homogenizer beads and ice-cold RNA Lysis/Binding buffer were added to each sample. Tissues were homogenized using an Omni Bead Ruptor (Omni International) for two cycles of 30 sec disruption until no large tissue chunks were observed. Samples were then centrifuged at 10,000xg for 20 min and the supernatant was collected for RNA extraction. Total RNA was isolated using the mirVana RNA isolation kit (#AM1561, Invitrogen) according to the manufacturer’s instructions, with on-column DNAse treatment (#12185010, Invitrogen) performed during Wash I. After RNA concentrations were determined using a NanoDrop ND1000 spectrophotometer, RNA samples were stored at −80°C.

### RNA sequencing (RNA-seq) and bioinformatics analysis

RNA-seq was done as previously described [20]. Briefly, total RNA was isolated from the dorsal horn of the lumbar spinal cord using the miRVana kit (Applied Biosystems). RNA concentration and integrity were assessed with the Agilent RNA 6000 Nano kit on a Bioanalyzer 2100 (Agilent Technologies). Libraries were generated by reverse transcription to double-stranded cDNA, followed by end repair, 3′ adenylation, adaptor ligation, purification, and PCR amplification. Sequencing was carried out on the BGISEQ-500 platform to obtain ∼30 million 50 bp single-end reads per sample. Raw reads were filtered using SOAPnuke to remove low-quality reads, adaptor sequences, and reads with ambiguous bases. Clean reads were assembled into unigenes, annotated, and analyzed for expression levels and SNPs. Reads were mapped to the reference transcriptome with HISAT [31].

For the microglia RNAseq experiment, the samples were collected in two batches and sequencing was conducted by Azenta Life Sciences. To minimize technical variation, batch correction was applied to raw read counts using ComBat-seq [32]. Low-abundance transcripts (TPM < 1 across all samples) were excluded from further analysis. Data is visualized using principal component analysis and hierarchical clustering with Pearson correlation distance metric. Differentially expressed genes (DEGs) were identified from normalized log2-TPM values using a two-tailed t-test (p ≤ 0.01) with a fold-change cutoff of ≥ 2. Adjusted p values are reported using Benjamini and Hochberg false discovery rate multiple testing correction. DEGs were submitted to enrichr [33], to determine the biological annotations and pathways they are involved in. Specifically, we tested for enrichment of terms available in the ChEA, Reactome, GO biological process, transcription factor perturbations, Wiki pathway, and KEGG pathway libraries. Significant enrichments were determined with p ≤ 0.01 and at least two DEGs annotated.

### Chromatin immunoprecipitation (ChIP) and bioinformatics analysis

We conducted ChIP-seq using protocols adapted from Covaris truChIP chromatin shearing kit tissue SDS and Millipore magna ChIP G tissue kits as described before [29] with an H3K4me1 antibody (Abcam ab8895) on spinal microglia 14 days after sEV injection from the two biological replicates per condition. Each replicate was generated by pooling 6-8 mice. ChIP-seq library preparation and sequencing were performed by Azenta Life Sciences. Libraries were generated from ChIP DNA using Azenta’s Illumina ChIP DNA Library Preparation workflow. Sequencing was carried out on an Illumina platform using a 2 × 150 bp paired-end configuration with single indexing, yielding approximately 350 million raw paired-end reads per lane (∼105 GB). A minimum quality threshold of ≥80% bases ≥Q30 was achieved for all libraries. Azenta provided raw FASTQ files along with initial quality reports, adapter trimming, read mapping, and peak-calling outputs. For downstream analysis, raw sequencing reads were processed using a standardized computational pipeline. Sequencing adapters and low-quality bases were trimmed with Trimmomatic [34], and the resulting reads were aligned to the mm10 reference genome using Bowtie2 [35]. Alignments were filtered with samtools [36] to retain primary concordant reads with a mapping quality ≥30. Peak calling was performed using MACS2 [37], and peaks overlapping genome-specific blacklist regions were removed. Valid peaks across samples were merged; to ensure high stringency, only peaks consistently identified in both biological replicates were retained for further analysis.

### Statistical analysis

Data analysis was performed with GraphPad Prism 10.2.0. Behavioral data were analyzed using repeated measures two-way analysis of variance (ANOVA), followed by Bonferroni *post-hoc* test for multiple comparisons and presented as mean ± standard error of the mean (SEM).

## Results

### sEV characterization

Typical diameter of sEVs range from 50-150 nm [38] and our NTA analysis with standard beads of 70, 125, 200, and 400 nm in diameter, showed that the sEVs had an average diameter of 149.2 ± 4.9 nm (Fig. 1A). We previously reported that sEVs from RAW 264.7 cells had a mean diameter of 113.6 ± 7.9 nm [20]. Western blotting of sEVs confirmed the expression of the exosome marker protein CD81 but not the negative control protein calnexin (Fig.1B). We also characterized sEV markers using a MACSPlex flow cytometry bead-based capture assay and detected the common sEV markers including tetraspanins CD81, CD9, and CD63. sEV composition reflects characteristics of their parent cell [39] and we observed expression of monocytic/macrophage markers in RAW 264.7 derived sEVs including CD115 (CSF1R) and CD86 (Fig. 1C). These results indicate that sEVs isolated meet the MISEV guidelines [27] for purity.

**Fig. 1.**
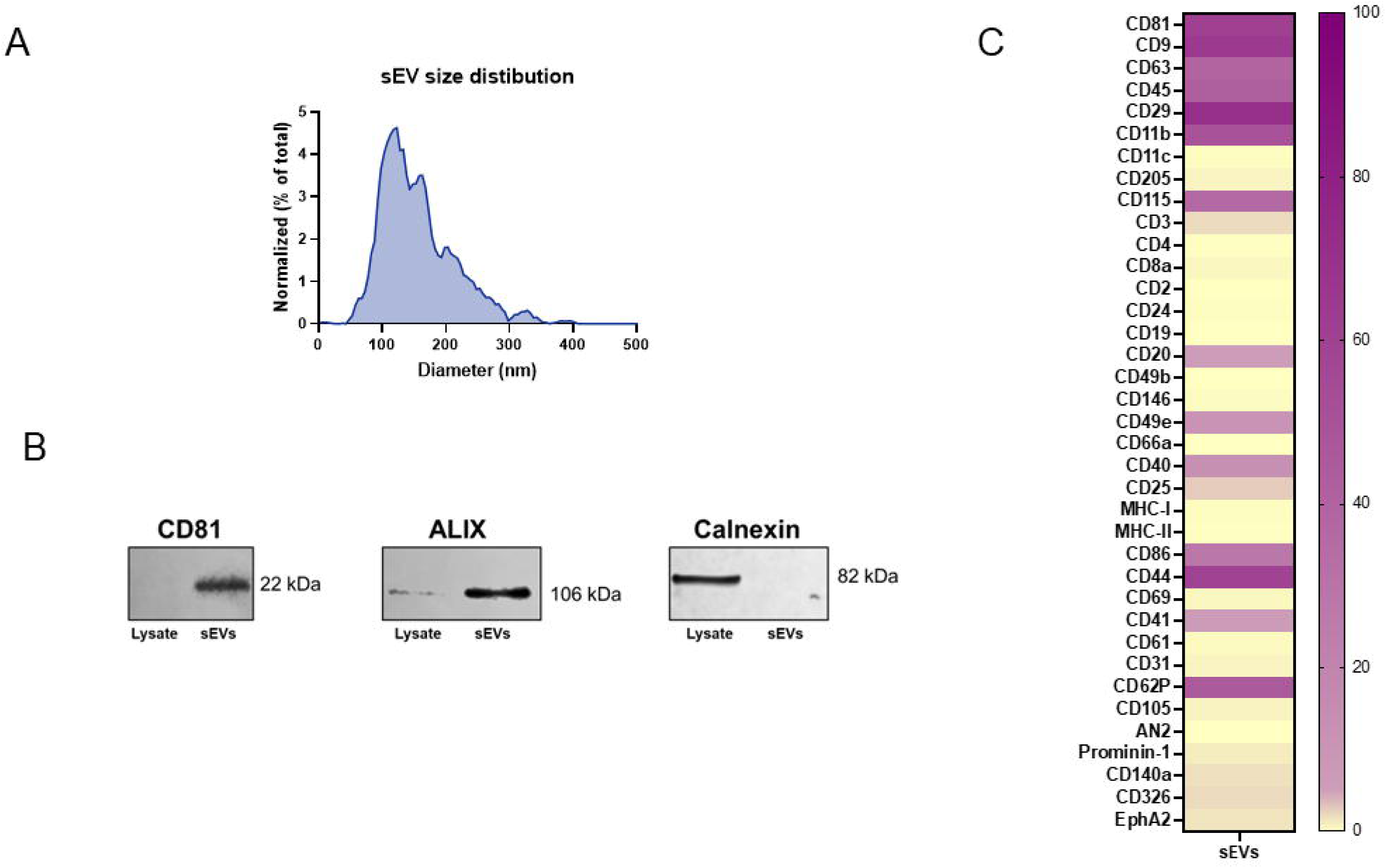
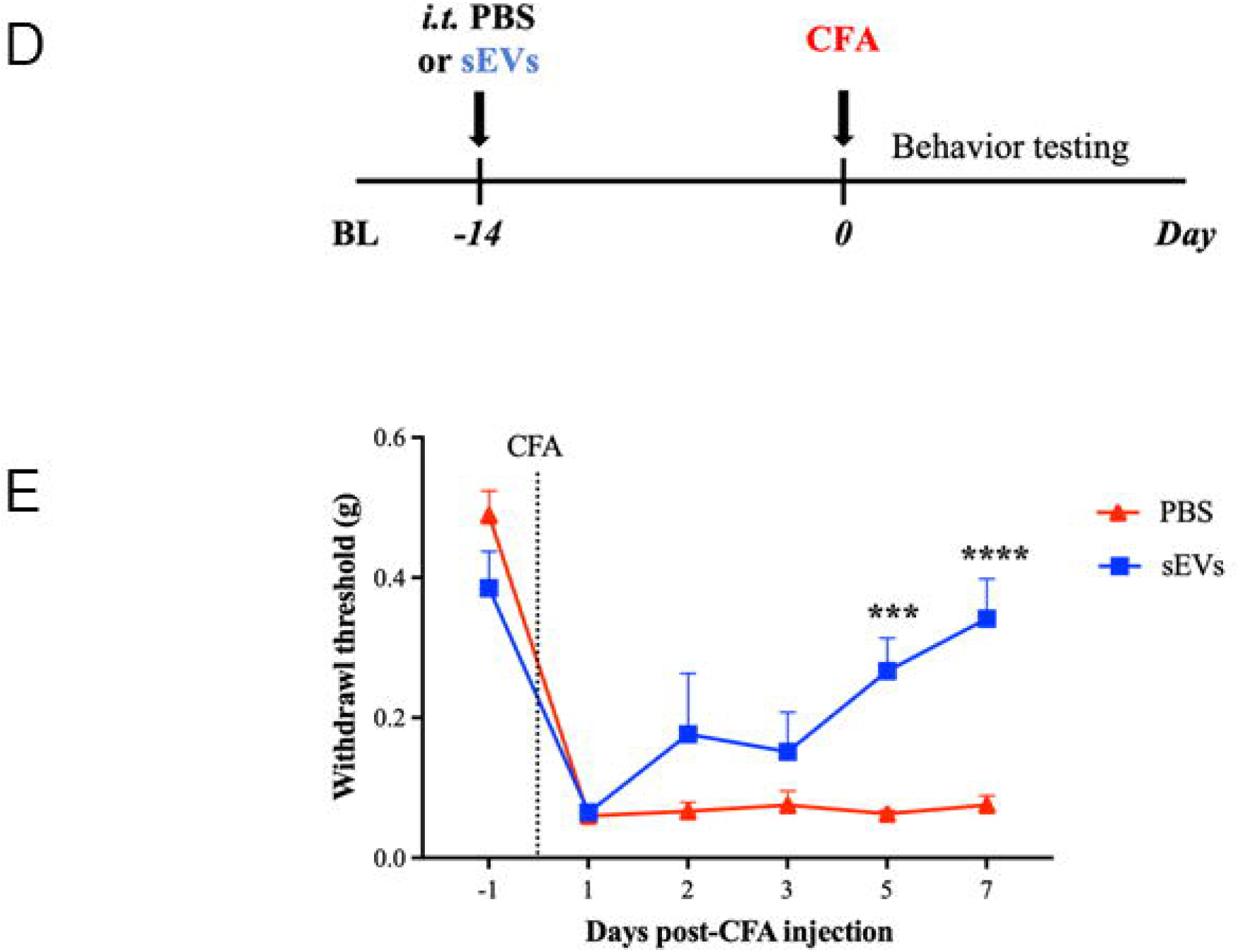
Confirmation of purity and *in vivo* efficacy of small extracellular vesicles (sEVs) from RAW 264.7 cells. **A)** Nanoparticle tracking analysis (NTA) showing an average diameter of 149.2 ± 4.9 nm in sEVs isolated from the conditioned medium of RAW 264.7 cells. **B)** Western blotting showing the presence of EV markers CD81 and ALIX in sEV preparations, with the negative marker calnexin only expressed in cell lysate. **C)** Bead-based flow cytometry confirmed the presence of sEV markers such as tetraspanins CD81, CD9, and CD63 and monocytic/macrophage markers in RAW 264.7-derived sEVs including CD115 (CSF1R) and CD86. Percent weighted expression calculated from fraction of positive beads and normalized mean fluorescent intensity. Purple indicates high expression and yellow indicates low expression. **D)** Schematic representation of prophylactic injection of sEVs in a mouse model of CFA-induced inflammatory pain. **E)** Prophylactic intrathecal administration of 1µg sEVs 14 days before CFA injection in the hind paw induces faster resolution of mechanical sensitivity. Macrophage sEVs significantly alleviate inflammatory pain starting from day 5. Data shown as mean ± SEM, n=9 for PBS and n=6 for sEVs. ***, ****p < 0.001, 0.0001 repeated measures two-way ANOVA with Bonferroni test. BL, baseline. CFA, complete Freund’s adjuvant. sEVs, small extracellular vesicles.

### Prophylactic intrathecal sEV administration attenuates inflammatory pain

In our previous studies, prophylactic intrathecal administration of 1 µg of macrophage-derived sEVs two weeks prior effectively attenuated CFA-induced inflammatory pain in male C57BL/6J mice, starting from two days post-CFA injection [20]. To independently confirm and replicate these effects in CFA model mice we followed the same paradigm (Fig. 1D). Our data showed that macrophage-derived sEVs significantly reduced inflammatory pain hypersensitivity on days 5 and 7 after CFA injection (Fig. 1E). Overall, our behavioral results demonstrate that macrophage-derived sEVs accelerate recovery from CFA-induced pain.

### Microglial depletion and repopulation using PLX5622 does not influence normal sensitivity in mice

To study the role of microglia in sEV-mediated pain resolution, we used a selective colony stimulating factor 1 receptor (CSF1R) inhibitor PLX5622. CSF1R serves as a key regulator of microglia and macrophages and is essential for myeloid cell survival [40]. Numerous studies have validated the use of CSF1R inhibitors in microglial ablation [41]. Interestingly, microglia can be fully repopulated in the CNS within 7 days after the cessation of CSF1R inhibition [42]. This is particularly advantageous for our study, as the absence of microglia may impact the development and maintenance of inflammatory pain [24, 43], making it difficult to interpret behavioral data. To induce microglial ablation, we fed the mice with PLX5622 (1,200 ppm) in AIN-76A diet (PLX). AIN-76A diet (control diet or CD) was used as the control. We examined microglial depletion in lumbar spinal cord in mice fed with PLX for 7 and 11 days. IHC was performed on lumbar segments (L4-L5) collected from these mice using the microglial-specific marker Iba1 (shown in red) and counterstained with DAPI (blue) to identify microglia (Fig. 2). Confocal images confirmed that microglia were ablated in spinal cord 7 days after PLX diet, with further depletion over an additional 4 days of PLX diet (Fig. 2A). To ensure complete ablation of microglia at the time of sEV injection, we extended the feeding schedule of PLX diet to 14 days. After depletion, the PLX diet was replaced with CD for a minimum of 10 days, allowing sufficient time for microglial repopulation to occur prior to CFA injection. Microglia were repopulated in the lumbar spinal cord 10 days after cessation of the PLX5622 diet (Fig. 2B).

**Fig. 2.**
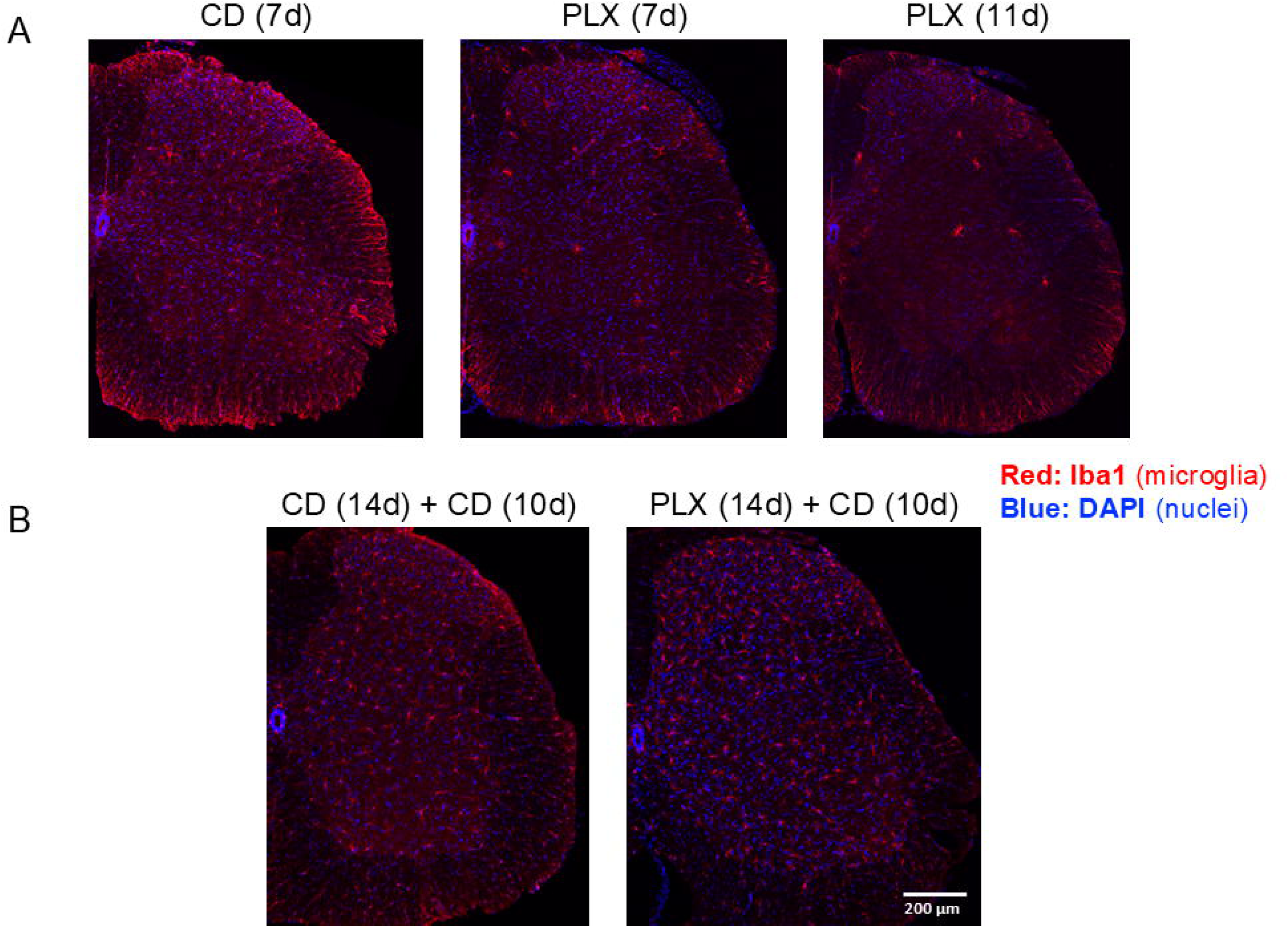
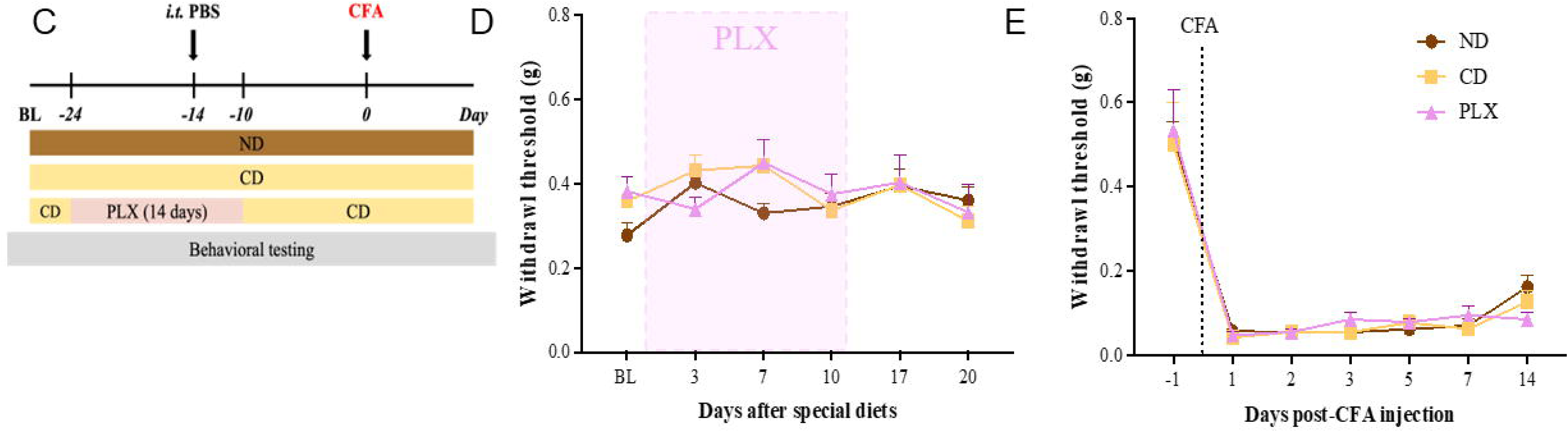
Depletion and repopulation of microglia in lumbar spinal cord and its effect on CFA-induced inflammatory pain. **A)** IHC images of lumbar spinal cord (L4-L5) confirm depletion of microglia 7 days and 11 days after PLX5622 diet. **B)** IHC also confirms microglial repopulation in L4–L5 spinal cord when mice were fed for 14 days with PLX5622 diet, followed by 10 days of control diet. Iba1 shown in red was used as microglial marker and counterstained with DAPI (blue). Scale bar = 200 µm. CD, control AIN-76A diet. PLX, AIN-76A diet containing PLX5622. **C)** Schematic of experiment to test if repopulated microglia alter CFA-induced hypersensitivity. In PLX group, mice were fed for 14 days with PLX5622 diet (before and after intrathecal injection of PBS), followed by 10 days of control diet. Comparison of the effects of three types of diets on normal sensitivity **D)** and CFA-induced pain hypersensitivity **E)** No differences in mechanical allodynia were observed among these three diets before and after CFA injections (repeated measures two-way ANOVA, Bonferroni post-test). Data presented as mean ± SEM (n=6 for ND; n=7 for CD and PLX).

To investigate whether microglial depletion and repopulation affect basal sensitivity and CFA-induced pain hypersensitivity, we conducted behavioral testing using von Frey filaments on mice fed with different diets before and after CFA injection. We also included a group of mice that were fed with a normal diet (ND) provided by our animal facility for comparison with CD and PLX diets. The schematic of experimental design is shown in Fig. 2C. Mice on CD and PLX diets showed no significant alteration in basal pain thresholds (Fig. 2D). Moreover, mice on all diets demonstrated similar pain sensitivity in response to CFA injection (Fig. 2E). We conclude that microglial ablation and repopulation does not influence normal mechanical sensitivity in mice.

### Microglia are indispensable for macrophage sEV-induced prophylaxis

We next investigated whether prophylactic sEVs could still confer early pain resolution when microglia were absent during sEV administration. After 10 days on PLX diet, mice were injected *i.t.* with 1 µg of sEVs or PBS. Microglial depletion continued for an additional 4 days, after which PLX diet was replaced with CD for the rest of the study (Fig. 3A). Control mice were fed with CD throughout the experiment. Prophylactic sEVs provided protection against CFA-induced pain in CD-fed mice, starting from day 5 and continuing until day 14 (Fig. 3B). However, the early resolution of mechanical hypersensitivity induced by sEVs was completely abolished in PLX-fed mice (Fig. 3B), indicating that microglia at the time of sEV administration were necessary for sEVs to promote faster resolution of inflammatory pain in CFA model mice.

**Fig. 3.**
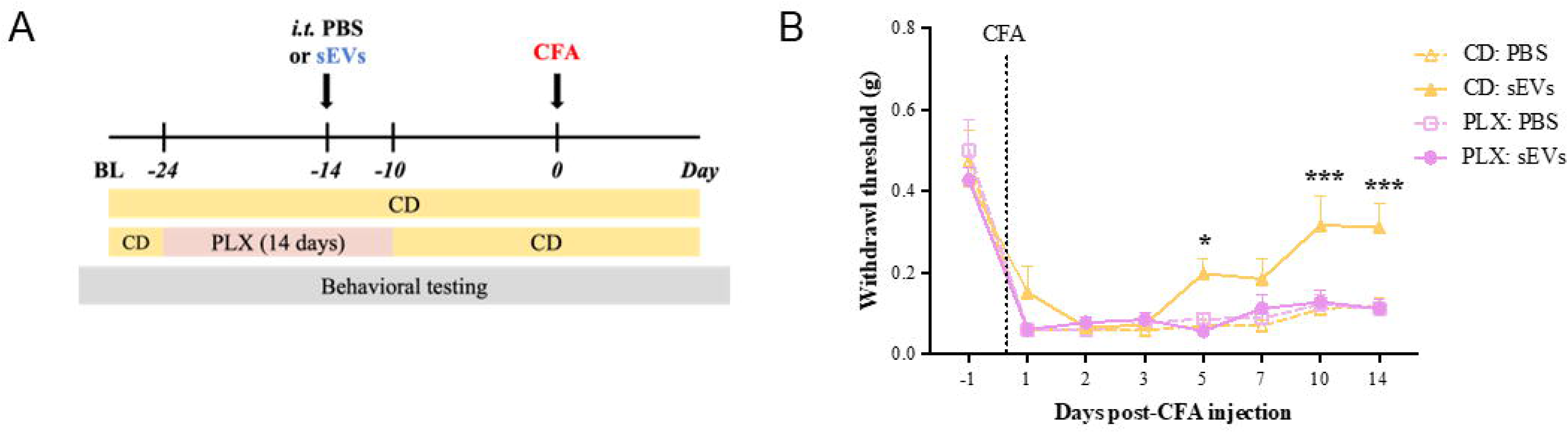
Spinal cord microglia are essential for macrophage sEV-induced prophylaxis. **A)** Experimental design. Mice were fed with PLX5622 diet (PLX) or AIN-76A control diet for 14 days (10 days before and 4 days after) sEV injection. **B)** sEV-induced prophylaxis was diminished in the mice fed with PLX diet during sEV injection. von Frey test for mechanical sensitivity showed that pain attenuation conferred by sEVs is abolished in PLX-fed mice. All mice received CFA. Data presented as the mean ± SEM (n=7-8; \**p* < 0.05, \*\*\**p* < 0.001 vs. PLX: sEVs, repeated measures two-way ANOVA with Bonferroni post-hoc test). BL, baseline. CFA, complete Freund’s adjuvant. sEVs, small extracellular vesicles. CD, control AIN-76A diet. PLX, AIN-76A diet containing PLX5622.

### sEVs induce gene expression changes in lumbar spinal cord 7 days after *i.t.* injection

To evaluate the transcriptional effects of sEV treatment on spinal cord tissue, we performed bulk RNA-seq on lumbar spinal cord segments (L4-L5) collected 7 days after intrathecal injection of sEVs or PBS. We identified 3 upregulated and 34 downregulated genes in sEV-treated spinal cords compared to PBS controls. The top 10 DEGs are listed in Table **1** and the whole list is shown in Supplementary Table 1. Notably, several small nucleolar RNAs (e.g., Snord116, Snord35a, and Snord89) and microRNAs (e.g., mir-9-2, mir-453) were among the most significantly altered, suggesting that sEVs may influence non-coding RNA regulation in the spinal cord. Three genes, somatostatin receptor 4, carbonic anhydrase 1, and regulatory subunit of type II PKA R-subunit (RIIa) domain containing 1 have been implicated in regulating pain [44–46]. To further investigate the biological relevance of the transcriptional changes, we performed gene set enrichment analysis (GSEA), which revealed significant associations with oxidative stress and neuroendocrine signaling. Genes altered by sEV treatment were enriched in NFE2L2 knockout signatures, suggesting modulation of antioxidant pathways. Additional enrichment was observed in pathways related to peptide hormone secretion and neuropeptide signaling, indicating potential effects on hormonal and synaptic regulation within the spinal cord (Supplementary Table 1).

**Table 1.**
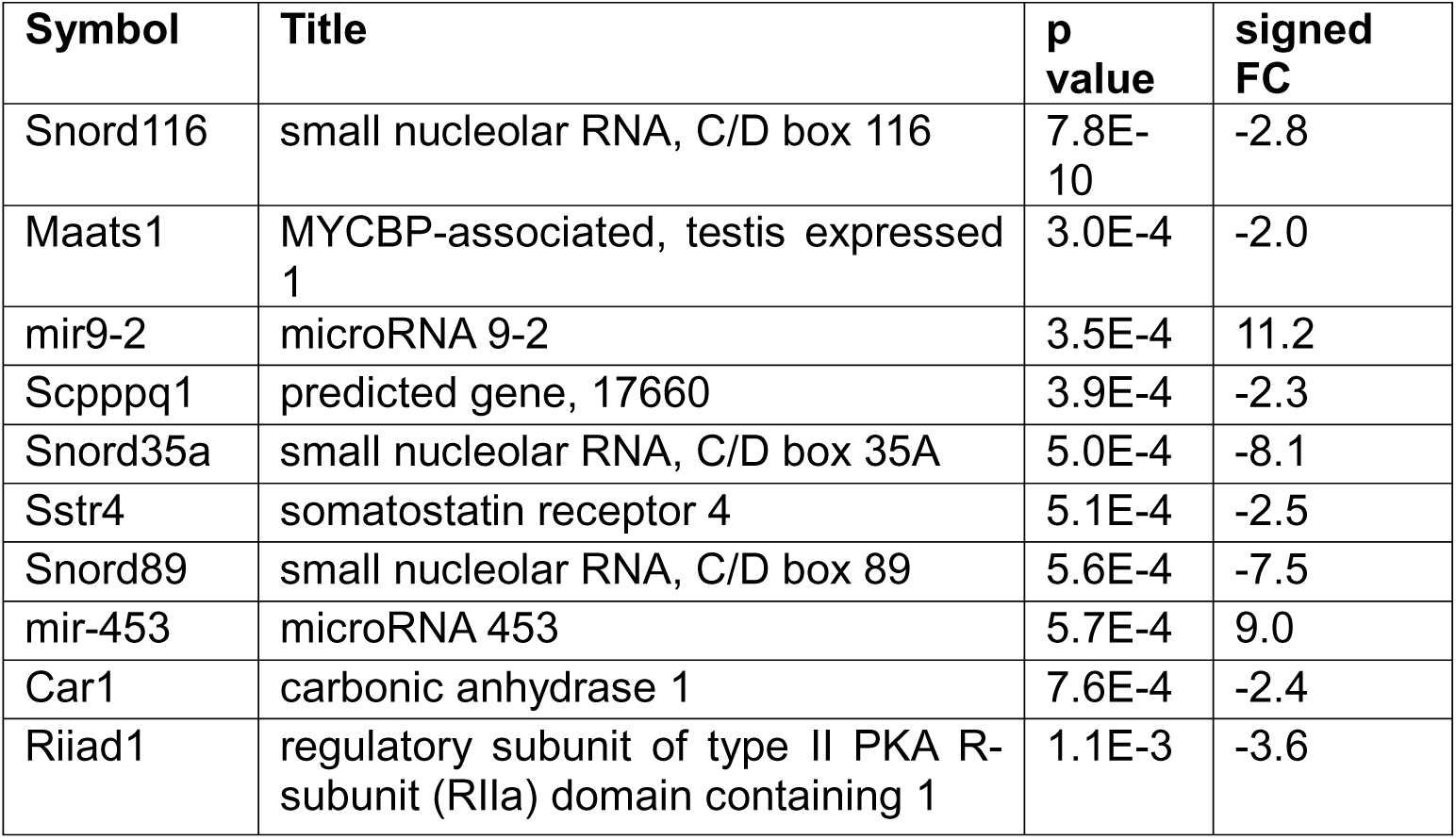
The top 10 differentially expressed genes in lumbar spinal cord tissue 7 days after sEV treatment. Genes were selected based on p-value < 0.01 and absolute fold change > 2. Signed fold change indicates direction and magnitude of expression change relative to PBS-treated controls. List of all DEGs and the enrichment results are available in Supplementary Table 1.

### sEV treatment alters microglial gene expression in lumbar spinal cord at 14 days

Next we used isolated microglia from the whole spinal cord of mice. Mice that received PBS were used as control. To obtain sufficient RNA yield for RNA-seq, spinal cord samples were pooled. Each sequencing replicate consisted of a pool of 6-8 mice. We used two MACS and two FACS-sorted CD11b+ microglia. The gating strategy is shown in Supplementary Fig. 1. One sEV sample (sEV2) was determined to be an outlier via PCA and clustering and was excluded from the analysis (Supplementary Fig. 2A and B). Differential expression analysis of the remaining samples identified 24 upregulated and 26 downregulated genes in sEV-treated microglia compared to PBS treated control microglia. Most of these DEGs were predicted or unannotated genes; top 10 annotated DEGs are listed in Table 2 and the full list is included in Supplementary Table 2. Gene set enrichment analysis revealed that the DEGs were significantly associated with immune and inflammatory signaling pathways. Notably, enriched terms included cytokine-cytokine receptor interaction, chemokine signaling, and malaria-related pathways, driven by genes such as CD40LG, CCL8, XCL1, and CCR1L1. These findings suggest that sEVs modulate microglial immune activity and chemokine-mediated communication in the spinal cord.

**Table 2.** The top 10 differentially expressed genes (excluding predicted or unannotated genes) in spinal cord microglia isolated 14 days after sEV treatment. Genes were selected based on p-value < 0.01 and absolute fold change > 2. Signed fold change indicates direction and magnitude of expression change relative to PBS-treated controls. List of all DEGs and the enrichment results are available in Supplementary Table 2.

| Symbol | Title | p value | signed FC |
| --- | --- | --- | --- |
| Igkj5 | immunoglobulin kappa joining 5 | 7.4E-5 | 4.4 |
| Ccl8 | C-C motif chemokine ligand 8 | 1.3E-3 | 2.3 |
| Mettl5os | methyltransferase 5, opposite strand | 1.6E-3 | -2.3 |
| Mir713 | microRNA 713 | 2.3E-3 | 3.5 |
| Snora62 | small nucleolar RNA, H/ACA box 62 | 2.7E-3 | 4.1 |
| Hoga1 | 4-hydroxy-2-oxoglutarate aldolase 1 | 2.8E-3 | -2.3 |
| Cd40lg | CD40 ligand | 2.9E-3 | 2.0 |
| Snora17 | small nucleolar RNA, H/ACA box 17 | 2.9E-3 | -2.4 |
| 1700001D01Rik | RIKEN cDNA 1700001D01 gene | 3.3E-3 | 3.6 |
| Ccr1l1 | C-C motif chemokine receptor 1 like 1 | 4.1E-3 | -2.3 |

### sEVs alter H3K4me1 enrichment in spinal microglia

To determine if the alterations in H3K4me1 enrichment following sEV treatment contribute to the resolution of CFA-induced inflammatory pain, we conducted chromatin immunoprecipitation followed by sequencing (ChIP-seq) with an H3K4me1 antibody on MACS-sorted CD11b^+^ microglia from ipsilateral lumbar L1-L6 spinal cord of CFA model mice that prophylactically received PBS or sEVs 14 days prior to CFA injection. Each replicate consisted of a pool of 6-8 mice and from the two biological samples per condition, we identified 12 merged annotated peaks in spinal microglia from PBS-treated mice and 85 merged annotated peaks in sEV-treated samples (Supplementary Table 3). There were no overlapping peaks between the two conditions. In the PBS-treated sample, most peaks (83.33%) were annotated to the promoter regions within 1 kb of the transcription start site (TSS), while the remainder were located in the distal intergenic regions (> 1.5 kb from TSS). Similarly, in the sEV-treated group, the majority of the peaks were also annotated to the promoter regions, and the average signal profiles indicated that the binding of H3K4me1 was enriched at TSSs. To explore the functional relevance of these sEV-specific H3K4me1-enriched regions, we performed GSEA using the genes associated with peaks present only in the sEV-treated group (Supplementary Table 3). The analysis revealed strong enrichment for targets of Polycomb group proteins and chromatin regulators, including SUZ12, MTF2, JARID2, and EZH2, suggesting that sEVs may influence epigenetic remodeling in spinal microglia. Additionally, enriched pathways included neuronal signaling, chemokine signaling, and synaptic regulation, with genes such as Gata2, Cacna1e, Doc2b, Pde10a, and Bcl11b contributing to these signatures. These findings support the hypothesis that sEVs modulate microglial function through epigenetic mechanisms that prime transcriptional programs relevant to neuronal and immune signaling.

### Cross-experiment comparisons reveal distinct gene and pathway responses to sEV treatment, with transcription factor-mediated links

To assess the relationship between transcriptional and epigenetic responses to sEV treatment, we compared results across bulk RNA-seq (7 days post sEV treatment lumbar spinal cord), purified microglial RNA-seq (14 days post sEV treatment), and microglia ChIP-seq (14 days post sEV treatment) experiments. As shown in Fig. 4A, there were no overlapping genes among the datasets, indicating that sEVs modulate distinct gene sets depending on cell type and time point. Similarly, gene set enrichment analysis revealed no shared pathways across experiments (Fig. 4B), suggesting functional divergence in sEV-induced responses. To identify potential regulatory connections, we focused on transcription factors identified as either differentially expressed or enriched in one dataset and mapped their known targets in other datasets. Notably, only transcription factors differentially expressed or enriched in the ChIP-seq dataset were found to target the differentially expressed genes in other datasets. These relationships, visualized in Fig. 5, highlight transcription factor-mediated links across experiments and suggest that sEVs orchestrate coordinated but compartmentalized regulatory programs in spinal cord microglia and tissue.

**Fig. 4.**
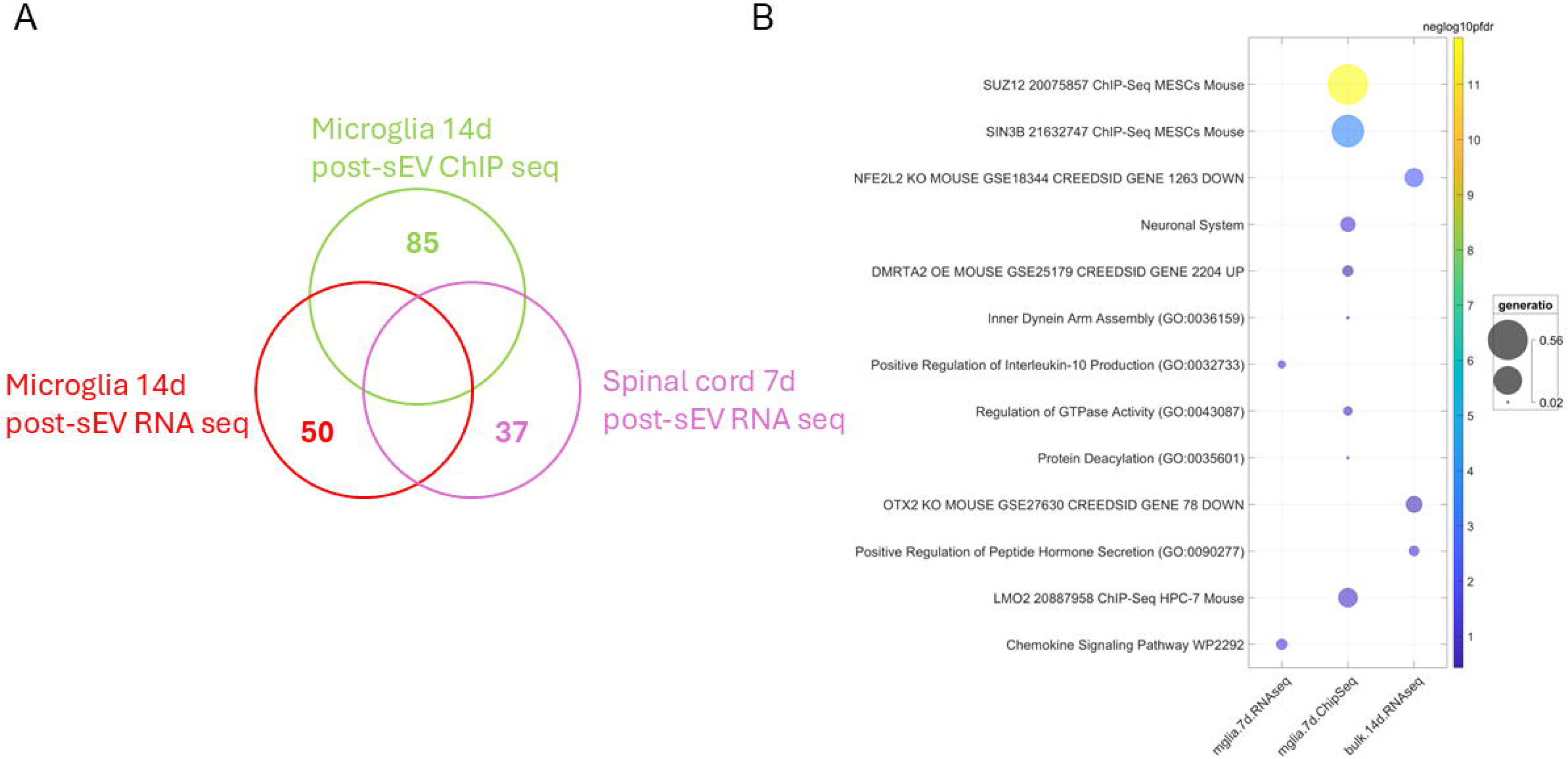
Cross-experiment comparisons reveal distinct gene and pathway responses to sEV treatment. **A)** Venn diagram showing the uniqueness and lack of overlap of genes identified across bulk RNA-seq, microglia RNA-seq, and ChIP-seq experiments. Each set represents genes significantly altered in spinal cord tissue or microglia after sEV treatment. **B)** Bubble chart of enriched gene sets from different experiments. Bubble size is proportional to the fraction of enriched DEGs annotated to that term (gene ratio) and the color reflects statistical significance of enrichment, scaled by the negative log10 of the false discovery rate (FDR). To reduce redundancy, gene sets sharing more than 50% of their genes were filtered, retaining only the most statistically significant term.

**Fig. 5.**
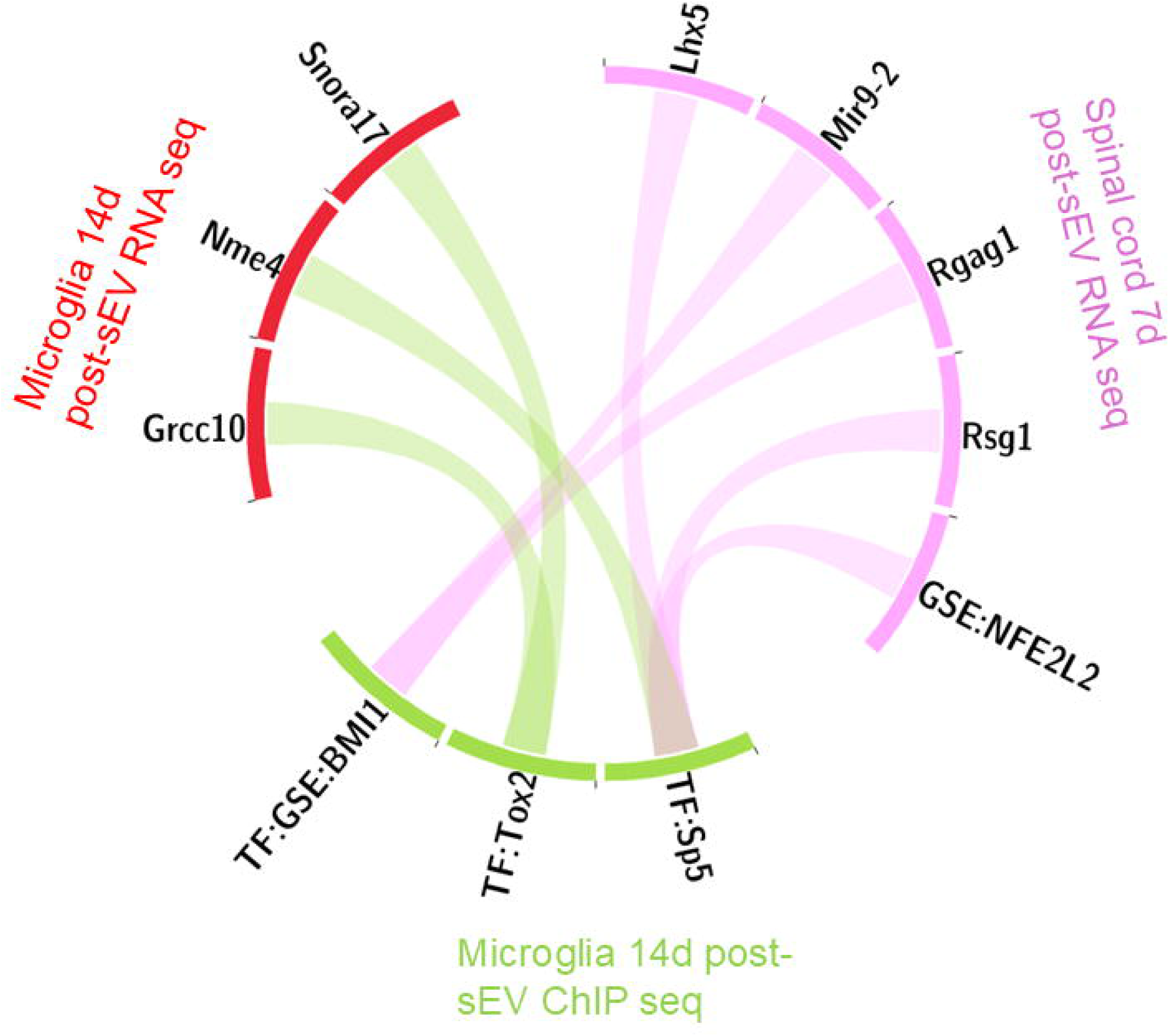
Circos diagram illustrating transcription factor-mediated regulatory connections across datasets. Transcription factors (TF) that are either differentially expressed or are enriched in the ChIP-seq dataset were mapped to their known target genes identified in RNA-seq experiments. There were no such connections originating from the RNA-seq experiments.

### Methyltransferase activity of SETD7 is essential for prophylaxis conferred by sEVs

To determine the necessity of H3K4me1 deposition for sEVs to confer faster resolution of pain hypersensitivity, we treated the mice with PFI-2, a selective methyltransferase inhibitor which blocks the methyltransferase activity of SETD7 [47] (Fig. 6A). PFI-2 was given via intrathecal catheters immediately after sEV delivery and repeated every day for an additional four days. Detailed experimental design and grouping are listed in Supplementary Table 4. Mice treated with PFI-2 showed no difference in mechanical thresholds post-CFA as compared to vehicle, indicating PFI-2 treatment alone did not affect the induction of the CFA model (Fig. 6B). Mice treated with sEVs followed by PFI-2 however showed significantly lower mechanical thresholds at days 7, 10, and 14 post-CFA than those treated with sEVs and vehicle. However, it is worth noting that multiple prophylactic injections of DMSO in the control groups (VEH: PBS, VEH: sEVs, and PFI-2: PBS), also facilitated faster recovery from CFA-induced pain as DMSO has anti-inflammatory effects [48]. Despite the confounding effects of DMSO, VEH:sEVs were still able to provide significant pain protection at day 14 post-CFA compared to VEH: PBS.

**Fig. 6.**
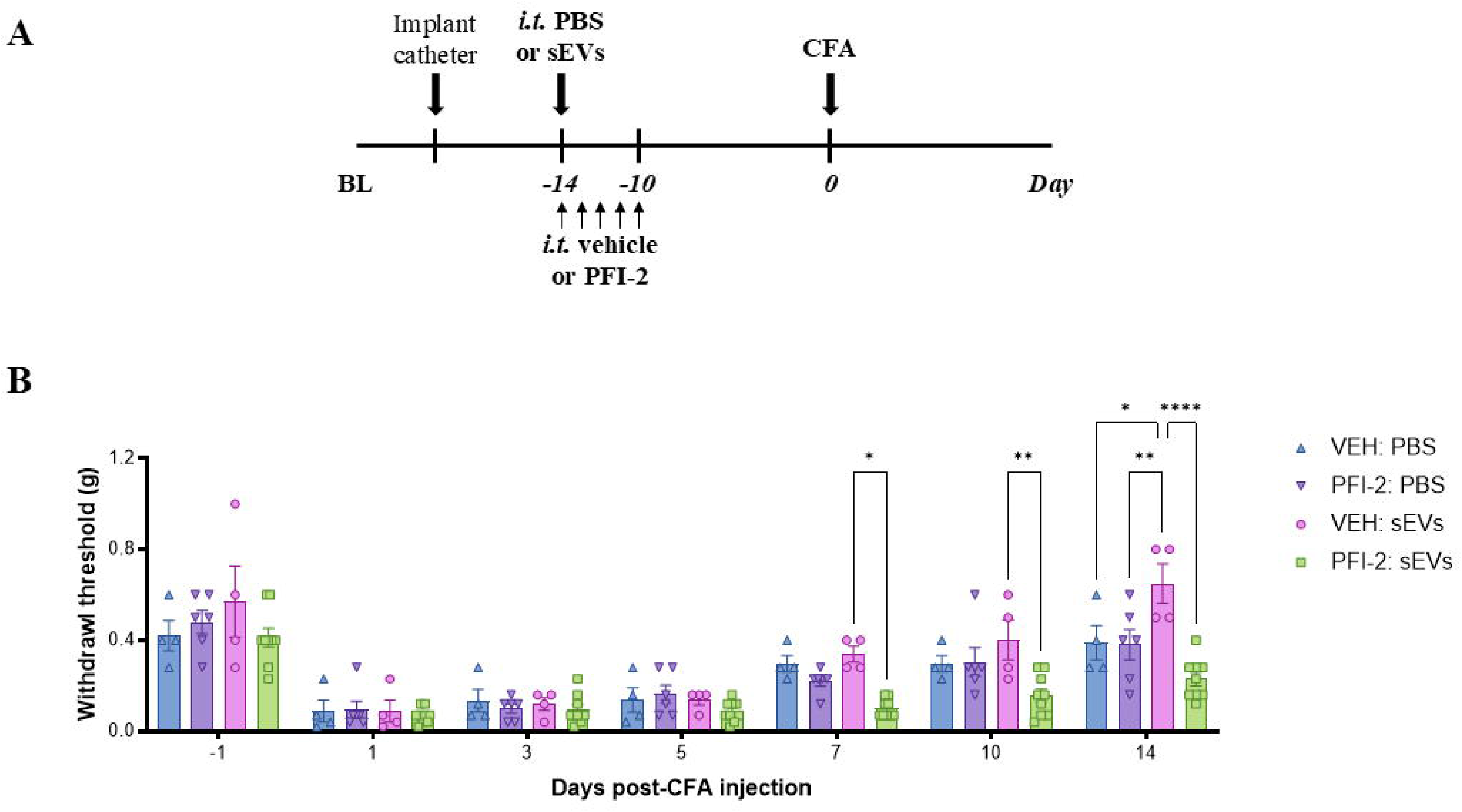
Inhibition of SETD7 mono-methylation activity blocks sEV-induced pain attenuation. **A)** Schematic representations of the experimental design. Mice were implanted with intrathecal catheters for repeated intrathecal injections. On the first injection day, mice received either sEVs or PBS. Additionally, all mice were administered PFI-2 or DMSO (VEH) for five consecutive days. Two weeks after sEV or PBS injection, CFA was injected into the hind paw. **B)** and **C)** Mechanical sensitivity tested in mice with distinct intrathecal injections. Data shown as mean ± SEM (n=4-9) and analyzed by repeated measures two-way ANOVA with Bonferroni multiple comparison test. *, **, \*\*\*\**p* < 0.05, 0.01, 0.0001, respectively. BL, baseline. sEVs, small extracellular vesicles. CFA, complete Freund’s adjuvant. VEH, vehicle.

## Discussion

The concept of innate immune memory is largely unexplored in the context of sEV biology [49] and pain. Based on our observation that 14-day prophylactic administration of sEVs from RAW 264.7 macrophage leads to faster resolution of mechanical and thermal hypersensitivity induced by CFA, we investigated whether these macrophage-derived sEVs confer long-lasting memory to induce protection against a subsequent painful insult. Recent findings suggest that microglia, the tissue-resident long-lived macrophages and primary innate immune cells of the CNS, can develop this form of immune memory [14, 50]. Our previous work illustrated that spinal microglia take up macrophage-derived sEVs when injected intrathecally [20, 51]. Thus, we hypothesized that microglia in the spinal cord are involved in sEV-induced prophylaxis. To test this, we used PLX5622, a highly selective CSF1R inhibitor, to ablate microglia [40].

PLX5622 can be formulated into a chow diet, which minimizes the stress on mice from multiple injections. We first confirmed microglial ablation in the spinal cord with the PLX5622 diet. IHC results showed that microglia were ablated in the lumbar (L4–L5) spinal cord 7 days after initiation of PLX5622 diet compared to control with an additional qualitative decrease in microglia numbers by day 11. This is in line with previous studies that a 14-day PLX5622 treatment is required to induce a stable microglial depletion in CNS [52, 53]. Therefore, we adhered to a 14-day PLX5622 treatment for subsequent studies. Interestingly, cessation of PLX5622 leads to the repopulation of microglia [42]. Since microglia play a significant role in regulating inflammatory pain [24], we switched mice back to a control diet for at least 10 days following the PLX5622 diet to allow for maximal microglia repopulation in the spinal cord which we also confirmed by IHC. To rule out any confounding factors from the PLX5622 diet, we also confirmed that neither the PLX5622 nor control diet affected basal sensation. To ensure microglia depletion at the time of intrathecal PBS/sEV injection but sufficient time for microglial repopulation prior to CFA we injected PBS/sEVs intrathecally 10 days after initiation of PLX5622 treatment.

Behavioral data demonstrated that sEV-mediated pain prophylaxis was completely lost in mice fed with PLX5622 diet during sEV delivery. This finding suggests the crucial role of microglia in the prophylactic effects of sEVs in the CFA model. However, although PLX5622 is widely used to eliminate microglia in the CNS, and depleted cells are generally identified as microglia, it should be noted that CSF1R is also expressed in other myeloid cells, including macrophages and myeloid progenitors [54]. Even short-term PLX5622 treatment can induce persistent changes in myeloid and lymphoid cells [55]. In addition, it is possible that sEVs indirectly modulate microglia by acting on other cell types within the spinal cord, which then impart epigenetic changes in microglia during the early phase (4 days after sEV injection when PLX5622 was discontinued). The timing of depletion and repopulation poses difficulties for conducting short-term depletion/repopulation in the late phase after administering sEVs. Moreover, a recent study has shown that depleting microglia during stimulation can lead to morphologically distinct microglia with altered transcriptomes [56]. We cannot rule out that administrating sEVs during microglial depletion may influence the repopulated microglia, although we saw no effect of microglia depletion alone on mechanical thresholds post-CFA induction.

One characteristic of innate immune memory is that innate cells, once triggered by a primary stimulus for a transient response, typically return to their basal transcription level, leaving the changes at the epigenetic level and/or in cellular metabolism. These modifications are responsible for subsequent transcriptional alterations upon later or secondary triggers to form a heightened and rapid response [57]. Active enhancers are identified by enrichments of both H3K27ac and H3K4me1, while H3K4me1 alone indicates an enhancer in an inactive, poised state. Immune stimulation results in *de novo* deposition of H3K27ac at distal regulatory regions and leads to increased H3K4me1 marks, which persist even after the primary stimulus and H3K27ac marks are removed [12]. Enhancers carrying only H3K4me1 facilitate a more rapid histone acetylation and gene induction upon subsequent stimulation. Thus, the retention of H3K4me1 contributes to an epigenomic memory of the initial stimulation [15]. We first investigated transcriptomic changes in PBS- and sEV-treated mice by performing bulk RNA-seq on lumbar spinal cord collected 7 days after sEV injection. We observed 3 upregulated and 34 downregulated genes. Several small nucleolar RNAs and miRNAs (miR-9-2, miR-453) were among the most altered, suggesting sEV-mediated modulation of noncoding RNA expressions. We then performed RNA-seq on CD11b^+^ microglia isolated from the whole spinal cord and identified 24 upregulated and 26 downregulated genes in sEV-treated microglia compared to control. GSEA showed that the DEGs were associated with immune and inflammatory signaling pathways suggesting that sEVs modulate microglial immune activity and chemokine-mediated communication in the spinal cord.

As gene expression changes in sEV treated mice at day 7 and 14 post-CFA were limited compared to controls, the factors that contribute to subsequent protection may be due to regulatory events at the epigenetic level. Enhancers are enriched for DNA motifs recognized by TFs [58] and serve as platforms that integrate transcription factor binding and, through chromatin looping, communicate with gene promoters to modulate transcriptional output [59]. RNA polymerase II gene transcription burst frequency is primarily encoded in enhancers and burst size in core promoters [60]. Enhancers can thus fine-tune gene expression by bridging transcription factors and the RNA polymerase II complex, thereby linking regulatory activity to gene transcription. Our data suggest that sEVs modulate microglial function through epigenetic priming of neuronal and immune signaling programs. It is important to note that the current analysis is unable to identify H3K4me1 peaks in enhancer regions. Enhancers can be located up to 1 million bp (Mbp) away from the TSS. Therefore a combination of other histone marks, such as H3K4me3 and H3K27ac [61], or more sophisticated algorithms [62] are necessary to determine if putative enhancer regions with long-lasting H3K4me1 depositions are present.

To examine the interplay between transcriptional and epigenetic responses to sEV treatment, we integrated RNA-seq data (lumbar spinal cord 7 days), microglia (14 days), and microglia ChIP-seq (14 days). There were no overlapping genes detected across these datasets, and gene set enrichment analysis revealed no shared pathways, indicating that sEVs influence distinct gene networks depending on cell type and temporal context. To explore potential regulatory connections, we focused on transcription factors differentially expressed or enriched within each dataset and mapped their known targets across experiments. Notably, only transcription factors with peaks identified in the ChIP-seq dataset were predicted to regulate differentially expressed genes in the RNA-seq datasets. These relationships suggest that sEVs may exert coordinated yet compartmentalized control of transcriptional programs in spinal microglia and surrounding tissue. The absence of overlap between bulk spinal cord (RNA-seq) and microglia-specific (RNA-seq/ChIP-seq) datasets or shared pathway enrichment may suggest temporal differences but is also in alignment with emerging evidence that microglial epigenomic remodeling is frequently uncoupled from steady-state transcriptional output. This disconnect is consistent with prior work showing that microglia-specific transcriptional signatures are often masked in heterogeneous cells [63, 64] Additionally, microglia undergo extensive enhancer reorganization in response to environmental cues or injury without corresponding immediate gene expression changes. Microglial enhancer landscapes are highly plastic and respond to cytokines, neuronal signals, and environmental perturbations, yet many new or primed enhancers remain transcriptionally silent until secondary stimulation [65, 66]. Similarly, microglia exposed to chronic stimuli or peripheral inflammation can acquire latent or trained epigenetic states that do not manifest in baseline transcriptomes but alter future responsiveness [12, 14]. Temporal dynamics are also critical as microglial transcription typically shows rapid, transient waves after injury or immune stimulation [63, 64], whereas enhancer remodeling marked by H3K4me1 can persist for weeks after initial activation. Together, these observations suggest that the sEV-induced microglial chromatin changes we observe at 14 days may represent a long-lasting regulatory state that does not necessarily coincide with stable transcriptional changes at the same time point or in the surrounding tissue environment.

We next tested if inhibiting the H3K4 methyltransferase SETD7 to prevent H3K4me1 deposition abolished sEV-induced pain attenuation. SETD7 has been identified for its role in mono-methylation of H3K4 [67, 68] in the context of innate immune memory. Increased expression of SETD7 in spinal microglia from mice with chronic constriction injury (CCI) correlated with elevated H3K4me1 levels [69]. We utilized the selective SETD7 inhibitor (*R*)-PFI-2 to inhibit SETD7 and reduce H3K4me1 deposition, following the same pharmacological approach that mitigated CCI-induced pain [69]. SETD7 inhibition for 5 consecutive days at the time of sEV administration significantly reduced the mechanical thresholds post-CFA as compared to vehicle treated mice, suggesting the importance of H3K4me1 deposition for mediating prophylactic sEV mediated pain resolution.

While these findings provide important insights, this study has limitations. Although we performed ChIP-seq for H3K4me1 to identify putative enhancer elements in spinal microglia following sEV treatment, we did not perform complementary ChIP-seq for H3K27ac. It is known that H3K4me1 alone marks primed enhancers, whereas H3K27ac is required to identify enhancers that are transcriptionally active or dynamically regulated in response to environmental cues [70]. Because active enhancers are typically defined by the combinatorial presence of H3K4me1 and H3K27ac, the absence of H3K27ac profiling limits our ability to distinguish poised from active enhancer states and to comprehensively map stimulus-responsive regulatory regions. Thus, incorporating H3K27ac ChIP-seq in future work will be essential for a more complete understanding of enhancer activation in microglia in response to sEVs. In addition, performing RNA-seq studies after CFA induction, and correlating them with ChIP-seq results would aid in confirming functional alterations in a secondary response for genes with altered H3K4me1 deposition. Another limitation is that only male mice were used in this study. Although a male-specific microglial contribution to neuropathic pain is reported [43], subsequent work has shown that this dichotomy is not universal. Several studies have reported microglial activation and functional involvement in females as well, depending on the pain model, injury type, and inflammatory context [71]. Microglia exhibit sex differences not only in pain mechanisms but also in their transcriptional and epigenomic landscapes. Prior studies have shown that male and female microglia differ in baseline chromatin accessibility, enhancer usage, and responsiveness to inflammatory stimuli, which can shape divergent pain pathways [72, 73]. Though sEVs are efficacious in both sexes, future work will determine whether sEV-mediated microglial epigenomic responses differ between males and females.

Overall, our findings showed that intrathecal injection of macrophage-derived sEVs required microglia to exert their prophylactic effects on accelerating inflammatory pain resolution. Inhibition of H3K4 mono-methyltransferase SETD7 during prophylactic sEV administration also prevented accelerated pain resolution. Given sEV induced alterations in H3K4me1 enrichment on specific gene loci in microglia, these changes may contribute to the prophylactic effects against CFA-induced pain. Although chronic pain is prevalent, a prophylactic strategy has not yet been tested in chronic pain models. Other prophylactic strategies that induce innate immune memory such as administration of live vaccines, induce protection against non-target diseases [74], which suggests that prophylactic macrophage sEVs could offer non-specific protection against subsequent pain insults. This opens the possibility for such prophylactic strategies to be extended to other pain and non-pain conditions and decrease opioid prescriptions. Our findings here provide a mechanistic basis that lays the foundation for these possibilities.

## References

1. Ji R-R, Chamessian A, Zhang Y-Q: Pain regulation by non-neuronal cells and inflammation. Science 2016, 354:572–577.

2. Dogan N, Wu W, Morrissey CS, Chen KB, Stonestrom A, Long M, Keller CA, Cheng Y, Jain D, Visel A, et al: Occupancy by key transcription factors is a more accurate predictor of enhancer activity than histone modifications or chromatin accessibility. Epigenetics Chromatin 2015, 8:16.

3. Ramaker RC, Hardigan AA, Goh ST, Partridge EC, Wold B, Cooper SJ, Myers RM: Dissecting the regulatory activity and sequence content of loci with exceptional numbers of transcription factor associations. Genome Res 2020, 30:939–950.

4. Cirovic B, de Bree LCJ, Groh L, Blok BA, Chan J, van der Velden W, Bremmers MEJ, van Crevel R, Handler K, Picelli S, et al: BCG Vaccination in Humans Elicits Trained Immunity via the Hematopoietic Progenitor Compartment. Cell Host Microbe 2020, 28:322–334 e325.

5. de Laval B, Maurizio J, Kandalla PK, Brisou G, Simonnet L, Huber C, Gimenez G, Matcovitch-Natan O, Reinhardt S, David E, et al: C/EBPbeta-Dependent Epigenetic Memory Induces Trained Immunity in Hematopoietic Stem Cells. Cell Stem Cell 2020, 26:793.

6. Litman GW, Anderson MK, Rast JP: Evolution of antigen binding receptors. Annu Rev Immunol 1999, 17:109–147.

7. Netea MG, Latz E, Mills KH, O’Neill LA: Innate immune memory: a paradigm shift in understanding host defense. Nat Immunol 2015, 16:675–679.

8. Netea MG, Joosten LA, Latz E, Mills KH, Natoli G, Stunnenberg HG, O’Neill LA, Xavier RJ: Trained immunity: A program of innate immune memory in health and disease. Science 2016, 352:aaf1098.

9. Netea MG, Quintin J, van der Meer JW: Trained immunity: a memory for innate host defense. Cell Host Microbe 2011, 9:355–361.

10. Lajqi T, Lang G-P, Haas F, Williams DL, Hudalla H, Bauer M, Groth M, Wetzker R, Bauer R: Memory-Like Inflammatory Responses of Microglia to Rising Doses of LPS: Key Role of PI3Kγ. Frontiers in Immunology 2019, Volume 10 - 2019.

11. Ostuni R, Piccolo V, Barozzi I, Polletti S, Termanini A, Bonifacio S, Curina A, Prosperini E, Ghisletti S, Natoli G: Latent enhancers activated by stimulation in differentiated cells. Cell 2013, 152:157–171.

12. Saeed S, Quintin J, Kerstens HHD, Rao NA, Aghajanirefah A, Matarese F, Cheng S-C, Ratter J, Berentsen K, van der Ent MA, et al: Epigenetic programming of monocyte-to-macrophage differentiation and trained innate immunity. Science 2014, 345:1251086.

13. Quintin J, Saeed S, Martens JHA, Giamarellos-Bourboulis EJ, Ifrim DC, Logie C, Jacobs L, Jansen T, Kullberg BJ, Wijmenga C, et al: Candida albicans infection affords protection against reinfection via functional reprogramming of monocytes. Cell Host Microbe 2012, 12:223–232.

14. Wendeln AC, Degenhardt K, Kaurani L, Gertig M, Ulas T, Jain G, Wagner J, Häsler LM, Wild K, Skodras A, et al: Innate immune memory in the brain shapes neurological disease hallmarks. Nature 2018, 556:332–338.

15. Bekkering S, Domínguez-Andrés J, Joosten LAB, Riksen NP, Netea MG: Trained Immunity: Reprogramming Innate Immunity in Health and Disease. Annu Rev Immunol 2021.

16. Groot Kormelink T, Mol S, de Jong EC, Wauben MHM: The role of extracellular vesicles when innate meets adaptive. Semin Immunopathol 2018, 40:439–452.

17. Kim SH, Kim S, Oligino TJ, Robbins PD: Effective treatment of established mouse collagen-induced arthritis by systemic administration of dendritic cells genetically modified to express FasL. Mol Ther 2002, 6:584–590.

18. McDonald MK, Tian Y, Qureshi RA, Gormley M, Ertel A, Gao R, Aradillas Lopez E, Alexander GM, Sacan A, Fortina P, Ajit SK: Functional significance of macrophage-derived exosomes in inflammation and pain. PAIN® 2014, 155:1527–1539.

19. Shiue SJ, Rau RH, Shiue HS, Hung YW, Li ZX, Yang KD, Cheng JK: Mesenchymal stem cell exosomes as a cell-free therapy for nerve injury-induced pain in rats. Pain 2019, 160:210–223.

20. Jean-Toussaint R, Lin Z, Tian Y, Gupta R, Pande R, Luo X, Hu H, Sacan A, Ajit SK: Therapeutic and prophylactic effects of macrophage-derived small extracellular vesicles in the attenuation of inflammatory pain. Brain Behav Immun 2021, 94:210–224.

21. Mianehsaz E, Mirzaei HR, Mahjoubin-Tehran M, Rezaee A, Sahebnasagh R, Pourhanifeh MH, Mirzaei H, Hamblin MR: Mesenchymal stem cell-derived exosomes: a new therapeutic approach to osteoarthritis? Stem Cell Res Ther 2019, 10:340.

22. Inoue K, Tsuda M: Microglia in neuropathic pain: cellular and molecular mechanisms and therapeutic potential. Nat Rev Neurosci 2018, 19:138–152.

23. Zhou LJ, Peng J, Xu YN, Zeng WJ, Zhang J, Wei X, Mai CL, Lin ZJ, Liu Y, Murugan M, et al: Microglia Are Indispensable for Synaptic Plasticity in the Spinal Dorsal Horn and Chronic Pain. Cell Rep 2019, 27:3844–3859 e3846.

24. Gu N, Yi MH, Murugan M, Xie M, Parusel S, Peng J, Eyo UB, Hunt CL, Dong H, Wu LJ: Spinal microglia contribute to sustained inflammatory pain via amplifying neuronal activity. Mol Brain 2022, 15:86.

25. Raghavendra V, Tanga FY, DeLeo JA: Complete Freunds adjuvant-induced peripheral inflammation evokes glial activation and proinflammatory cytokine expression in the CNS. Eur J Neurosci 2004, 20:467–473.

26. Chen G, Zhang YQ, Qadri YJ, Serhan CN, Ji RR: Microglia in Pain: Detrimental and Protective Roles in Pathogenesis and Resolution of Pain. Neuron 2018, 100:1292–1311.

27. Welsh JA, Goberdhan DCI, O’Driscoll L, Buzas EI, Blenkiron C, Bussolati B, Cai H, Di Vizio D, Driedonks TAP, Erdbrügger U, et al: Minimal information for studies of extracellular vesicles (MISEV2023): From basic to advanced approaches. J Extracell Vesicles 2024, 13:e12404.

28. Lin Z, Luo X, Wickman JR, Reddy D, DaCunza JT, Pande R, Tian Y, Kasimoglu EE, Triana V, Lee J, et al: Inflammatory pain resolution by mouse serum-derived small extracellular vesicles. *Brain*, Behavior, and Immunity 2025, 123:422–441.

29. Manners MT, Ertel A, Tian Y, Ajit SK: Genome-wide redistribution of MeCP2 in dorsal root ganglia after peripheral nerve injury. Epigenetics & Chromatin 2016, 9:23.

30. Pino PA, Cardona AE: Isolation of brain and spinal cord mononuclear cells using percoll gradients. J Vis Exp 2011.

31. Kim D, Langmead B, Salzberg SL: HISAT: a fast spliced aligner with low memory requirements. Nat Methods 2015, 12:357–360.

32. Zhang Y, Parmigiani G, Johnson WE: ComBat-seq: batch effect adjustment for RNA-seq count data. NAR Genomics and Bioinformatics 2020, 2.

33. Kuleshov MV, Jones MR, Rouillard AD, Fernandez NF, Duan Q, Wang Z, Koplev S, Jenkins SL, Jagodnik KM, Lachmann A, et al: Enrichr: a comprehensive gene set enrichment analysis web server 2016 update. Nucleic Acids Res 2016, 44:W90–97.

34. Bolger AM, Lohse M, Usadel B: Trimmomatic: a flexible trimmer for Illumina sequence data. Bioinformatics 2014, 30:2114–2120.

35. Langmead B, Salzberg SL: Fast gapped-read alignment with Bowtie 2. Nature Methods 2012, 9:357–359.

36. Danecek P, Bonfield JK, Liddle J, Marshall J, Ohan V, Pollard MO, Whitwham A, Keane T, McCarthy SA, Davies RM, Li H: Twelve years of SAMtools and BCFtools. Gigascience 2021, 10.

37. Gaspar JM: Improved peak-calling with MACS2. bioRxiv 2018:496521.

38. Kowal J, Tkach M, Thery C: Biogenesis and secretion of exosomes. Curr Opin Cell Biol 2014, 29:116–125.

39. Colombo M, Raposo G, Thery C: Biogenesis, secretion, and intercellular interactions of exosomes and other extracellular vesicles. Annu Rev Cell Dev Biol 2014, 30:255–289.

40. Elmore MR, Najafi AR, Koike MA, Dagher NN, Spangenberg EE, Rice RA, Kitazawa M, Matusow B, Nguyen H, West BL, Green KN: Colony-stimulating factor 1 receptor signaling is necessary for microglia viability, unmasking a microglia progenitor cell in the adult brain. Neuron 2014, 82:380–397.

41. Green KN, Crapser JD, Hohsfield LA: To Kill a Microglia: A Case for CSF1R Inhibitors. Trends Immunol 2020, 41:771–784.

42. Huang Y, Xu Z, Xiong S, Sun F, Qin G, Hu G, Wang J, Zhao L, Liang YX, Wu T, et al: Repopulated microglia are solely derived from the proliferation of residual microglia after acute depletion. Nat Neurosci 2018, 21:530–540.

43. Sorge RE, Mapplebeck JCS, Rosen S, Beggs S, Taves S, Alexander JK, Martin LJ, Austin J-S, Sotocinal SG, Chen D, et al: Different immune cells mediate mechanical pain hypersensitivity in male and female mice. Nat Neurosci 2015, 18:1081–1083.

44. Szolcsányi J, Pintér E, Helyes Z, Petho G: Inhibition of the function of TRPV1-expressing nociceptive sensory neurons by somatostatin 4 receptor agonism: mechanism and therapeutical implications. Curr Top Med Chem 2011, 11:2253–2263.

45. Supuran CT: Carbonic anhydrase inhibition and the management of neuropathic pain. Expert Rev Neurother 2016, 16:961–968.

46. Isensee J, Diskar M, Waldherr S, Buschow R, Hasenauer J, Prinz A, Allgöwer F, Herberg FW, Hucho T: Pain modulators regulate the dynamics of PKA-RII phosphorylation in subgroups of sensory neurons. Journal of Cell Science 2014, 127:216–229.

47. Barsyte-Lovejoy D, Li F, Oudhoff MJ, Tatlock JH, Dong A, Zeng H, Wu H, Freeman SA, Schapira M, Senisterra GA, et al: (R)-PFI-2 is a potent and selective inhibitor of SETD7 methyltransferase activity in cells. Proceedings of the National Academy of Sciences 2014, 111:12853–12858.

48. Elisia I, Nakamura H, Lam V, Hofs E, Cederberg R, Cait J, Hughes MR, Lee L, Jia W, Adomat HH, et al: DMSO Represses Inflammatory Cytokine Production from Human Blood Cells and Reduces Autoimmune Arthritis. PLoS One 2016, 11:e0152538.

49. Lajqi T, Köstlin-Gille N, Hillmer S, Braun M, Kranig SA, Dietz S, Krause C, Rühle J, Frommhold D, Pöschl J, et al: Gut Microbiota-Derived Small Extracellular Vesicles Endorse Memory-like Inflammatory Responses in Murine Neutrophils. Biomedicines 2022, 10.

50. Neher JJ, Cunningham C: Priming Microglia for Innate Immune Memory in the Brain. Trends Immunol 2019, 40:358–374.

51. Gupta R, Luo X, Lin Z, Tian Y, Ajit SK: Uptake of Fluorescent Labeled Small Extracellular Vesicles In Vitro and in Spinal Cord. J Vis Exp 2021, 171.

52. Brennan FH, Li Y, Wang C, Ma A, Guo Q, Li Y, Pukos N, Campbell WA, Witcher KG, Guan Z, et al: Microglia coordinate cellular interactions during spinal cord repair in mice. Nature Communications 2022, 13:4096.

53. Sawicki CM, Kim JK, Weber MD, Faw TD, McKim DB, Madalena KM, Lerch JK, Basso DM, Humeidan ML, Godbout JP, Sheridan JF: Microglia Promote Increased Pain Behavior through Enhanced Inflammation in the Spinal Cord during Repeated Social Defeat Stress. J Neurosci 2019, 39:1139–1149.

54. Hume DA, MacDonald KP: Therapeutic applications of macrophage colony-stimulating factor-1 (CSF-1) and antagonists of CSF-1 receptor (CSF-1R) signaling. Blood 2012, 119:1810–1820.

55. Lei F, Cui N, Zhou C, Chodosh J, Vavvas DG, Paschalis EI: CSF1R inhibition by a small-molecule inhibitor is not microglia specific; affecting hematopoiesis and the function of macrophages. Proc Natl Acad Sci U S A 2020, 117:23336–23338.

56. Donovan LJ, Bridges CM, Nippert AR, Wang M, Wu S, Forman TE, Haight ES, Huck NA, Bond SF, Jordan CE, et al: Repopulated spinal cord microglia exhibit a unique transcriptome and contribute to pain resolution. Cell Rep 2024, 43:113683.

57. Netea MG, Schlitzer A, Placek K, Joosten LAB, Schultze JL: Innate and Adaptive Immune Memory: an Evolutionary Continuum in the Host’s Response to Pathogens. Cell Host Microbe 2019, 25:13–26.

58. Heinz S, Romanoski CE, Benner C, Glass CK: The selection and function of cell type-specific enhancers. Nature Reviews Molecular Cell Biology 2015, 16:144–154.

59. Meng H, Bartholomew B: Emerging roles of transcriptional enhancers in chromatin looping and promoter-proximal pausing of RNA polymerase II. J Biol Chem 2018, 293:13786–13794.

60. Larsson AJM, Johnsson P, Hagemann-Jensen M, Hartmanis L, Faridani OR, Reinius B, Segerstolpe Å, Rivera CM, Ren B, Sandberg R: Genomic encoding of transcriptional burst kinetics. Nature 2019, 565:251–254.

61. Blinka S, Reimer MH, Jr., Pulakanti K, Pinello L, Yuan GC, Rao S: Identification of Transcribed Enhancers by Genome-Wide Chromatin Immunoprecipitation Sequencing. Methods Mol Biol 2017, 1468:91–109.

62. Butt AH, Alkhalifah T, Alturise F, Khan YD: A machine learning technique for identifying DNA enhancer regions utilizing CIS-regulatory element patterns. Sci Rep 2022, 12:15183.

63. Masuda T, Sankowski R, Staszewski O, Böttcher C, Amann L, Sagar, Scheiwe C, Nessler S, Kunz P, van Loo G, et al: Spatial and temporal heterogeneity of mouse and human microglia at single-cell resolution. Nature 2019, 566:388–392.

64. Hammond TR, Dufort C, Dissing-Olesen L, Giera S, Young A, Wysoker A, Walker AJ, Gergits F, Segel M, Nemesh J, et al: Single-Cell RNA Sequencing of Microglia throughout the Mouse Lifespan and in the Injured Brain Reveals Complex Cell-State Changes. Immunity 2019, 50:253–271.e256.

65. Gosselin D, Link VM, Romanoski CE, Fonseca GJ, Eichenfield DZ, Spann NJ, Stender JD, Chun HB, Garner H, Geissmann F, Glass CK: Environment drives selection and function of enhancers controlling tissue-specific macrophage identities. Cell 2014, 159:1327–1340.

66. Lavin Y, Winter D, Blecher-Gonen R, David E, Keren-Shaul H, Merad M, Jung S, Amit I: Tissue-resident macrophage enhancer landscapes are shaped by the local microenvironment. Cell 2014, 159:1312–1326.

67. Keating ST, Groh L, van der Heijden C, Rodriguez H, Dos Santos JC, Fanucchi S, Okabe J, Kaipananickal H, van Puffelen JH, Helder L, et al: The Set7 Lysine Methyltransferase Regulates Plasticity in Oxidative Phosphorylation Necessary for Trained Immunity Induced by beta-Glucan. Cell Rep 2020, 31:107548.

68. Benjaskulluecha S, Boonmee A, Pattarakankul T, Wongprom B, Klomsing J, Palaga T: Screening of compounds to identify novel epigenetic regulatory factors that affect innate immune memory in macrophages. Sci Rep 2022, 12:1912.

69. Shen Y, Ding Z, Ma S, Ding Z, Zhang Y, Zou Y, Xu F, Yang X, Schafer MKE, Guo Q, Huang C: SETD7 mediates spinal microgliosis and neuropathic pain in a rat model of peripheral nerve injury. Brain Behav Immun 2019, 82:382–395.

70. Creyghton MP, Cheng AW, Welstead GG, Kooistra T, Carey BW, Steine EJ, Hanna J, Lodato MA, Frampton GM, Sharp PA, et al: Histone H3K27ac separates active from poised enhancers and predicts developmental state. Proc Natl Acad Sci U S A 2010, 107:21931–21936.

71. Mogil JS, Parisien M, Esfahani SJ, Diatchenko L: Sex differences in mechanisms of pain hypersensitivity. Neurosci Biobehav Rev 2024, 163:105749.

72. Guneykaya D, Ivanov A, Hernandez DP, Haage V, Wojtas B, Meyer N, Maricos M, Jordan P, Buonfiglioli A, Gielniewski B, et al: Transcriptional and Translational Differences of Microglia from Male and Female Brains. Cell Reports 2018, 24:2773–2783.e2776.

73. Hanamsagar R, Alter MD, Block CS, Sullivan H, Bolton JL, Bilbo SD: Generation of a microglial developmental index in mice and in humans reveals a sex difference in maturation and immune reactivity. Glia 2017, 65:1504–1520.

74. Netea MG, Dominguez-Andres J, Barreiro LB, Chavakis T, Divangahi M, Fuchs E, Joosten LAB, van der Meer JWM, Mhlanga MM, Mulder WJM, et al: Defining trained immunity and its role in health and disease. Nat Rev Immunol 2020, 20:375–388.

