## Supplemental Table 4 for "Microglial epigenetic memory is associated with accelerated resolution of inflammatory pain induced by prophylactic macrophage-derived small extracellular vesicles"

| **Group** | **Day 1** | **Day 2-4** (10 µL) |
| --- | --- | --- |
| PBS/VEH | 5 µL 1% DMSO + 5 µL PBS | 0.1% DMSO |
| sEVs/VEH | 5 µL 1% DMSO + 5 µL sEVs (1 µg) | 0.1% DMSO |
| PBS/PFI-2 | 5 µL 20 µM PFI-2 + 5 µL PBS | 10 µM PFI-2 |
| sEVs/PFI-2 | 5 µL 20 µM PFI-2 + 5 µL sEVs (1 µg) | 10 µM PFI-2 |

**Supplementary Table 4**. Experimental design for studying effects of blocking H3Kme1 deposition by inhibiting SETD7 methyltransferase activity using PFI-2 in sEV-induced prophylaxis. DMSO at different concentrations were used as vehicle control. VEH, vehicle. sEVs, small extracellular vesicles.
