## Supplementary figures and images for "Microglial epigenetic memory is associated with accelerated resolution of inflammatory pain induced by prophylactic macrophage-derived small extracellular vesicles"

### Supplemental Figure 1

#### Slide 1
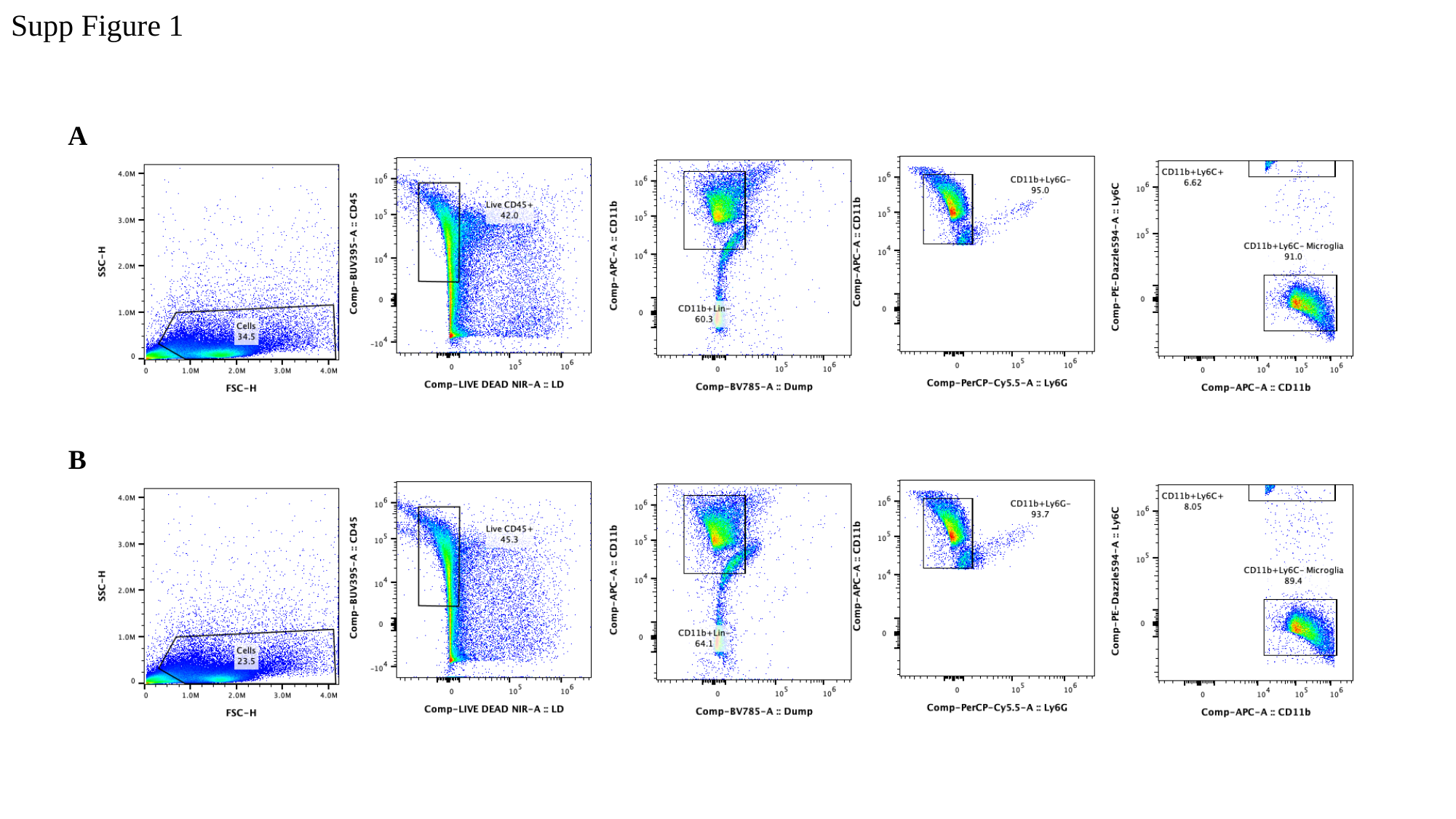

Supp Figure 1
A
B

### Supplemental Figure 2

#### Slide 1
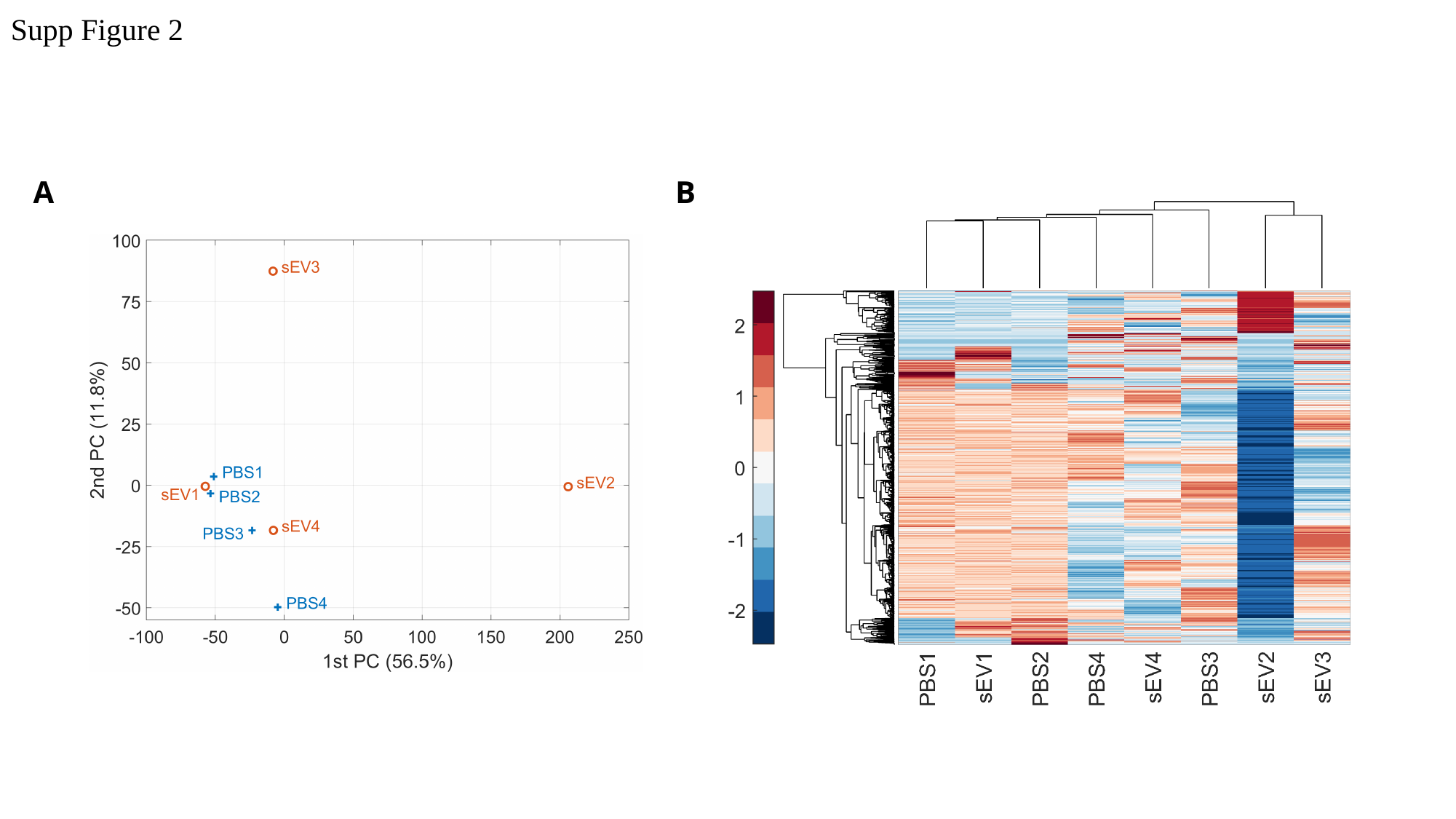

Supp Figure 2
A
B
